# ClpA- and ClpAP-Catalyzed Unfolding and Translocation are Differentially Coupled to ATP Binding

**DOI:** 10.1101/2025.08.12.669770

**Authors:** Liana Islam, Jaskamaljot Kaur Banwait, Aaron L. Lucius

## Abstract

ClpA is an ATP-dependent chaperone essential for protein quality control in *E. coli*. Upon ATP binding, ClpA forms hexameric rings capable of association with the tetradecameric ClpP protease. ClpA couples ATP binding and/or hydrolysis to the unfolding and translocation of protein substrates into the central cavity of ClpP for degradation. We previously developed a single-turnover stopped-flow method sensitive to ClpA-catalyzed translocation in the absence of ClpP-catalyzed proteolysis. This method was used on unstructured substrates so that the kinetics were reflective of translocation and not unfolding. We showed that at saturating [ATP], ClpA translocated at ∼20 aa s^-1^, with the kinetic step size, i.e., the average number of amino acids (aa) translocated between two rate-limiting steps being ∼14 aa step^-1^. Adding ClpP increased the rate to ∼36 aa s^-1^ and decreased the kinetic step-size to ∼5 aa step^-1^.

Here we apply this method to substrates containing folded Titin I27 domains. We report that at saturating [ATP], ClpA unfolded and translocated at ∼12 aa s^-1^, nearly half the rate of translocation alone. However, in the presence of ClpP, ClpA exhibited a rate of ∼40 aa s^-1^, representing no reduction in rate over translocation alone. Interestingly, unlike translocation alone, the kinetic step-size for unfolding and translocation was ∼29 aa step^-1^ for both ClpA and ClpAP. Examining the [ATP]-dependence of the unfolding reactions revealed that the increased kinetic step-size results from the averaging of a large unfolding step-size of ∼97 aa, representing cooperative unfolding of a single Titin I27 domain, followed by multiple smaller translocation steps on the newly unfolded chain. Moreover, just like translocation alone, the introduction of folds into the substrate results in different kinetics between ClpA and ClpAP. These observations further support a model where ClpP allosterically impacts ClpA-catalyzed processes.

**Significance:** ClpA is one of several AAA+ motors in *E. coli*. As part of the ATP-dependent protease ClpAP, it facilitates the removal of misfolded and properly folded proteins from the cell. Previously, we published the [ATP]-dependencies of kinetic parameters such as rate constants, kinetic step-sizes, and rates for ClpA- and ClpAP-catalyzed translocation. Here, for the first time, we make similar determinations for the unfolding and translocation cycle. We find both processes to be kinetically coupled to ATP binding, with unfolding being more sensitive to decreasing [ATP] compared to translocation. This coupling differs between ClpA and ClpAP. These findings reinforce the foundation for comparing how AAA+ motors respond to substrate folds, ATP levels, and allosteric regulation.

## Introduction

ATPases associated with diverse cellular activities (AAA+) are found in cells from all domains of life. AAA+ members are indispensable in protein quality control, the innate immune system, DNA replication, neuron polarization, mitochondrial division, and many other biological processes [1–5]. All members are molecular motors characterized by highly conserved ATPase domains.

ClpA is a AAA+ motor that couples energy from ATP binding and hydrolysis to unfolding and translocation of substrate proteins [6]. Upon binding ATP, ClpA assembles into hexameric rings capable of associating with the tetradecameric protease, ClpP, and with substrates targeted for unfolding [7, 8]. Associated with ClpP, ClpA catalyzes ATP-driven unfolding and translocation of the substrate into the proteolytic barrel of ClpP where the substrate is proteolyzed [9–11].

Previously, we developed a single-turnover fluorescence stopped-flow method to interrogate the mechanisms of ClpA and ClpAP-catalyzed protein translocation in the absence of unfolding [12, 13]. This was achieved by using substrates lacking any significant structure. In that work, we reported elementary rate constants, overall rates, and the first kinetic step-size for a protein unfoldase.

The kinetic step-size represents the average number of amino acids (aa) translocated between two rate-limiting steps. We found the kinetic step-size to be independent of [ATP] and ∼14 aa and ∼5 aa for ClpA and ClpAP, respectively, on substrates that lack significant fold. The lack of an [ATP] dependence suggests that the time courses are rate-limited by the same step within each repeating round of translocation [12–16].

From such experiments, the identity of the repeating step that occurs every 14 or 5 aa translocated is unknown. However, the observed rate constant was found to depend hyperbolically on [ATP]. This indicates that the observed repeating step immediately follows ATP binding. Thus, one can put constraints on step identity, e.g. it is neither ATP binding, nor ADP/Pi release. Thus, one is left with mechanical movement, ATP hydrolysis, or a slow conformational change as the identity of the observed step.

Single-molecule optical-trapping experiments have been used to examine ClpAP-catalyzed unfolding and translocation of tandem repeats of Titin I27 domains [17, 18]. ClpAP was shown to rapidly unfold a Titin I27 domain in a single step. Multiple translocation steps with a mechanical step-size of ∼5 amino acid step^-1^ follow on the newly unfolded polypeptide. This cycle repeats until all domains are unfolded, translocated, and degraded by ClpP. Notably, ClpAP spends more time translocating compared to the pre-unfolding dwell-time. This led to the conclusion that ClpAP-catalyzed protein unfolding is fast and translocation is rate-limiting [17].

Inspired by substrates used in the optical trapping experiments [19], and the RepA(1-70)-GFP substrates made by Wickner and Coworkers [20, 21], we constructed the RepA-Titin_X_ substrates, with repeating folded Titin I27 domains, see Figure 1 (A). Using these substrates, we developed a single-turnover stopped-flow method to examine unfolding catalyzed by ClpB, a AAA+ motor homologous to ClpA [22, 23].

**Figure 1:**
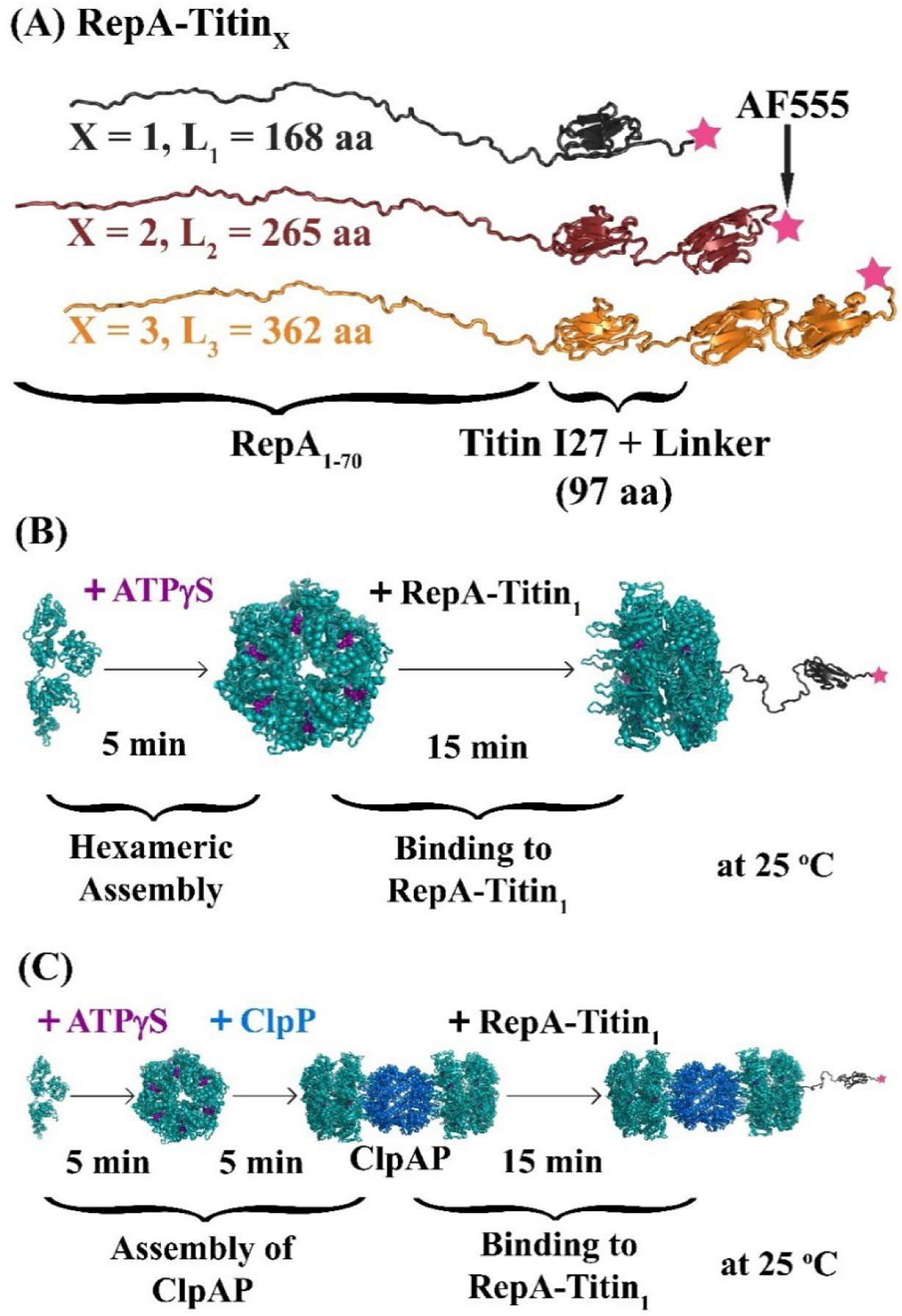
Substrates and incubation schemes. **(A)** Domain organization of RepA-Titin_1_ (black), RepA-Titin_2_ (brown), and RepA-Titin_3_ (orange). The N-terminal RepA_1-70_ is a binding sequence for ClpA and ClpAP. Tandem repeats of β-folded Titin I27 domains are separated by linkers. Each substrate is labeled with Alexa Fluor 555 (AF555) at the C-terminus (pink star). **(B)** Schematic for assembling hexameric ClpA (teal, nucleotide binding sites in deep purple) bound to RepA-Titin_X_. **(C)** Schematic for assembling ClpAP with hexameric ClpA binding tetradecameric ClpP (blue). ClpAP then binds RepA-Titin_X_.

Here we report single-turnover stopped-flow experiments to examine ClpA and ClpAP-catalyzed unfolding of RepA-Titin_X_. At saturating [ATP], ClpA and ClpAP exhibit the same kinetic step-size of ∼29 aa per rate limiting step. Interestingly, at reduced [ATP], the observed kinetic step-size for unfolding and translocation increases for both ClpA and ClpAP. This is in contrast with the [ATP]-independent kinetic step-sizes of ∼14 aa and ∼5 aa for ClpA and ClpAP-catalyzed polypeptide translocation, respectively [12, 13]. Here we show that the increased kinetic step-size for protein unfolding compared to translocation alone is the result of the averaging of a large unfolding step-size with a smaller translocation step-size, where the two processes exhibit differential coupling to ATP.

## Materials and Methods

### Reagents and Buffers

*E. coli* ClpA was purified as described [8]. All reported concentrations for ClpA are for monomers. We used ATP from ThermoFischer Scientific (Waltham, MA), and ATPγS from CalBiochem (La Jolla, CA). α-Casein (mixture of α-S1-Casein and α-S2-Casein) from Sigma-Aldrich (Darmstadt, Germany) was dissolved in 6 M guanidine hydrochloride with 20 mM HEPES (pH 7.5 at 25 °C) prior to dialysis into reaction conditions. RepA-Titin_X_ substrates were purified as described [22].

Buffers were made using reagent-grade chemicals and 18 MΩ water from a Purelab Ultra Genetic system (Evoqua, Warrendale, PA). Before experiments, reagents were dialyzed into buffer H300 containing 25 mM HEPES (pH 7.5 at 25 °C), 10 mM MgCl_2_, 300 mM NaCl, 2 mM 2-mercaptoethanol, and 10% (v/v) glycerol.

### RepA-TitinX Substrates

These were previously used to study ClpB-catalyzed unfolding and translocation [22, 23]. On the N-terminus, each substrate contains the RepA_1-70_ binding sequence for ClpA and ClpAP, see Figure 1 (A) [21, 24, 25]. This is followed by X = 1 to 3 repeats of β-folded Titin I27 domains [18]. The C-terminus has a cysteine labeled with Invitrogen Alexa Fluor 555 C2 Maleimide (AF555). To make sure that we are only labeling this C-terminal cysteine, all other cysteines within the Titin I27 domains were mutated to alanines. More details on these substrates have been previously published [22].

### Stopped-Flow Fluorescence Experiments

ClpA- and ClpAP-catalyzed protein unfolding and translocation were monitored using an SX.20 stopped-flow fluorometer (Applied Photophysics, Letherhead, UK). For ClpA-only experiments, ClpA was incubated with ATPγS for 5 min, then with RepA-Titin_X_ for 15 min at 25 °C, see Figure 1 (B). 1 μM ClpA, 150 μM ATPγS, and 100 nM RepA-Titin_X_ were loaded into Syringe 1, see Figure 2. For ClpAP-experiments, ClpA was incubated with ATPγS for 5 min, then with ClpP for 5 min, then with RepA-Titin_X_ for 15 min at 25 °C, see Figure 1 (C). 1 μM ClpA, 150 μM ATPγS, 1.2 μM ClpP, and 100 nM RepA-Titin_X_ were loaded into Syringe 1, see Figure 2. The incubation schemes were chosen based on previous research [7, 8, 12, 13, 21, 26–30]. Syringe 2 contained 20 μM α-Casein and varying ATP concentrations, also incubated at 25 °C for at least 10 min. The contents of both syringes were rapidly mixed 1:1 within 2 ms at 25 °C. Throughout the paper, [ATP] stands for the concentration of ATP post-mixing. Time courses were collected until the voltage level plateaued. Time courses were processed to represent the signal in terms of Relative Fluorescence Enhancement (RFE) as a function of time as previously described [22]. All time courses shown represent one individual determination. A total of three determinations were made at each [ATP].

**Figure 2:**
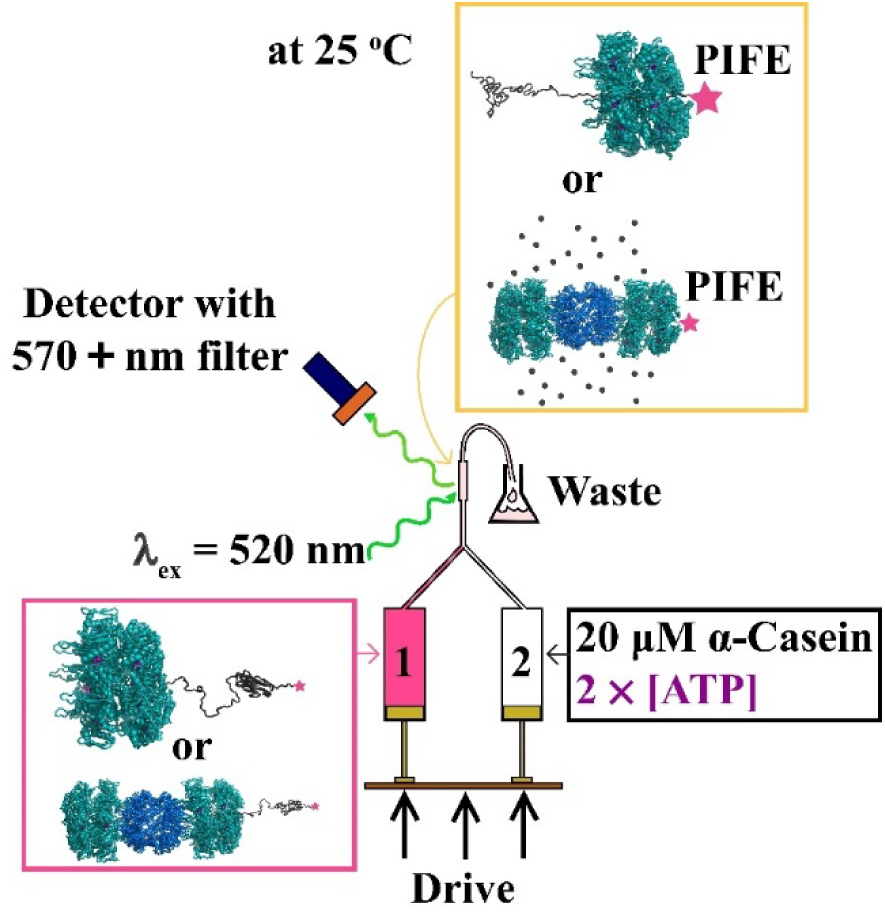
Single-turnover stopped-flow assay. Syringe 1 contains ClpA-bound (top of pink box) or ClpAP-bound (bottom of pink box) substrates. Syringe 2 contains the listed reagents. Before mixing the contents of Syringes 1 and 2, ClpA is at a distance greater than 3 nm from AF555 (pink star), and there is no Protein-Induced Fluorescence Enhancement (PIFE). After rapid mixing, AF555 in the solution was excited at 520 nm. Fluorescence emission from the mixing chamber was observed at 570 nm and above using a long-pass filter. We expect ClpA and ClpAP to arrive at AF555 following repeated ATP hydrolysis, resulting in PIFE, as shown in the yellow box.

### Fitting Time Courses

We solved the system of coupled differential equations describing the schemes using the method of Laplace transforms [31, 32]. The fitting functions thus obtained account for contributions from one or more intermediates, along with the unfolded substrate, RepA-Titin_XU_. The fitting process has been detailed in the supplementary section. We used the MATLAB (The MathWorks, Natick, MA) toolbox, MENOTR [33], to fit individual time courses to determine the kinetic parameters. Each parameter was averaged over three determinations.

### Modeling [ATP]-Dependencies

[ATP]-dependencies of kinetic parameters (velocities, rate constants, and overall rates) were analyzed by NLLS analysis using cooperative and noncooperative models. The cooperative model is defined by the Hill equation,

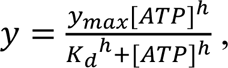

where *y* is the kinetic parameter, *y_max_* is the maximum value of the kinetic parameter at saturating [ATP], *K_d_* is the ATP dissociation equilibrium constant, and *h* is the Hill coefficient. The noncooperative model has *h* constrained to one.

## Results

### Substrate Length-Dependence

We hypothesized that folded domains would reduce the rates of ClpA- and ClpAP-catalyzed unfolding and translocation relative to translocation on unstructured substrates. To test this, we used the folded RepA-Titin_X_ substrates, see Figure 1 (A). ClpA requires nucleoside triphosphate binding to assemble into hexameric rings with substrate-binding and ClpP-binding activities. Thus, to induce assembly and substrate binding, ClpA or ClpAP was mixed with ATPγS and RepA-Titin_X_ following the incubation schemes in Figure 1 (B) or (C). The resulting complex was loaded into Syringe 1 of a stopped-flow fluorometer, see Figure 2. Syringe 2 was loaded with ATP and excess α-casein.

After rapid mixing of the contents of Syringes 1 and 2, the α-casein traps any unbound ClpA, thereby maintaining single-turnover conditions. We expected that upon binding ATP, ClpA or ClpAP would catalyze substrate unfolding, translocate the newly unfolded polypeptide chain, arrive at AF555 on the C-terminus to cause protein-induced fluorescence enhancement (PIFE), then slowly dissociate, resulting in loss of PIFE.

Figure 3 (A) shows a time course from ClpA-bound RepA-Titin_1_ at 5 mM ATP. The time course exhibits a lag, a peak, and a decay. We interpret the lag to represent the time over which unfolding and translocation occur. The peak, marked with the peak time, *t_peak,1_*, is interpreted as the arrival of ClpA within 3 nm of the C-terminal AF555 to cause PIFE. Because PIFE occurs when proteins are within 0-3 nm of the fluorophore [34–36], we interpret the decrease in Relative Fluorescence Enhancement (RFE) at the decay phase as dissociation of ClpA from unfolded RepA-Titin_1_.

**Figure 3:**
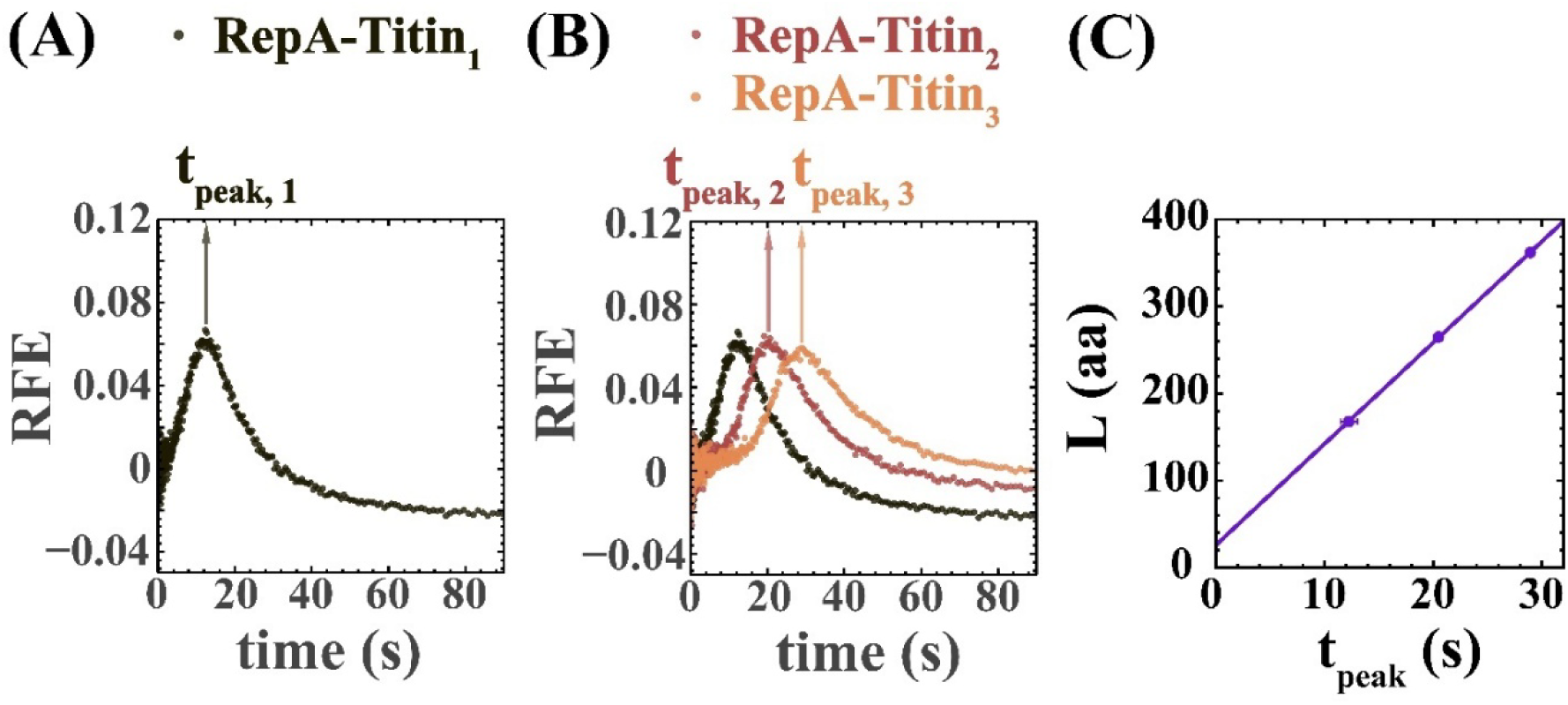
Peak time analysis for ClpA. **(A)** Representative time course for RepA-Titin_1_ at [ATP] = 5 mM displaying a lag, a peak, and a decay. **(B)** Time courses for all RepA-Titin_X_ substrates at the same [ATP]. **(C)** Substrate length, *L*, vs. peak time, *t_peak_*, reveals length dependence. Error bars: standard deviations (S.D.) in *t_peak_* for three replicates. Linear-least-squares analysis yields a slope (velocity), *v* = (11.6 ± 0.1) aa s^-1^ and *L_intercept_* = (26 ± 2) aa, with the errors representing standard errors (S.E.).

If the peak in Figure 3 (A) represents the arrival of ClpA to within 3 nm of the C-terminus, the peak should appear later for longer substrates. To test this, the experiments were repeated with RepA-Titin_2_ and RepA-Titin_3_. As expected, the peak emerges later for longer substrates, see Figure 3 (B). Substrate length, *L*, vs. *t_peak_*is linear, with the slope (velocity), *v* = (11.6 ± 0.1) aa s^-1^, and the intercept, *L_intercept_* = (26 ± 2) aa.

The impact of ClpP on ClpA-catalyzed protein unfolding was tested with similar experiments in the presence of ClpP. Although on a shorter time scale, the time courses exhibited the same overall shape as in Figure 3 (B) for ClpA alone, see Figure S1 (A)-(B). The time courses were subjected to the same “peak time analysis” and a velocity of *v* = (37.7 ± 0.4) aa s^-1^ with an intercept of *L_intercept_* = (14 ± 3) aa were determined, see Figure S1 (C). Thus, at 5 mM ATP, ClpAP unfolds proteins three times faster than ClpA, consistent with our previous observations that ClpAP is faster than ClpA for pure translocation, i.e., on unstructured substrates [12, 13, 37].

In both cases, a positive *L_intercept_* implies that some length of the substrate is already traversed before ATP addition. So far, *L_intercept_* is comparable to or less than the 24-25 aa substrate length seen in the axial channel of cryo-EM structures of ClpAP [29, 38]. Thus, *L_intercept_* may represent the length of substrate bound pre-translocation, i.e., the occluded or excluded length.

### [ATP]-Dependence

If the time courses are reporting on ATP-driven unfolding and translocation, they should also depend on [ATP]. To test this, we collected time courses by varying [ATP]. Each set of three time courses for the three RepA-Titin_X_ substrates were subjected to peak time analysis, see Figure S2 for *L* vs. *t_peak_*plots. The velocities, *v*, thus obtained were plotted against [ATP], see Figure 4 (A).

**Figure 4:**
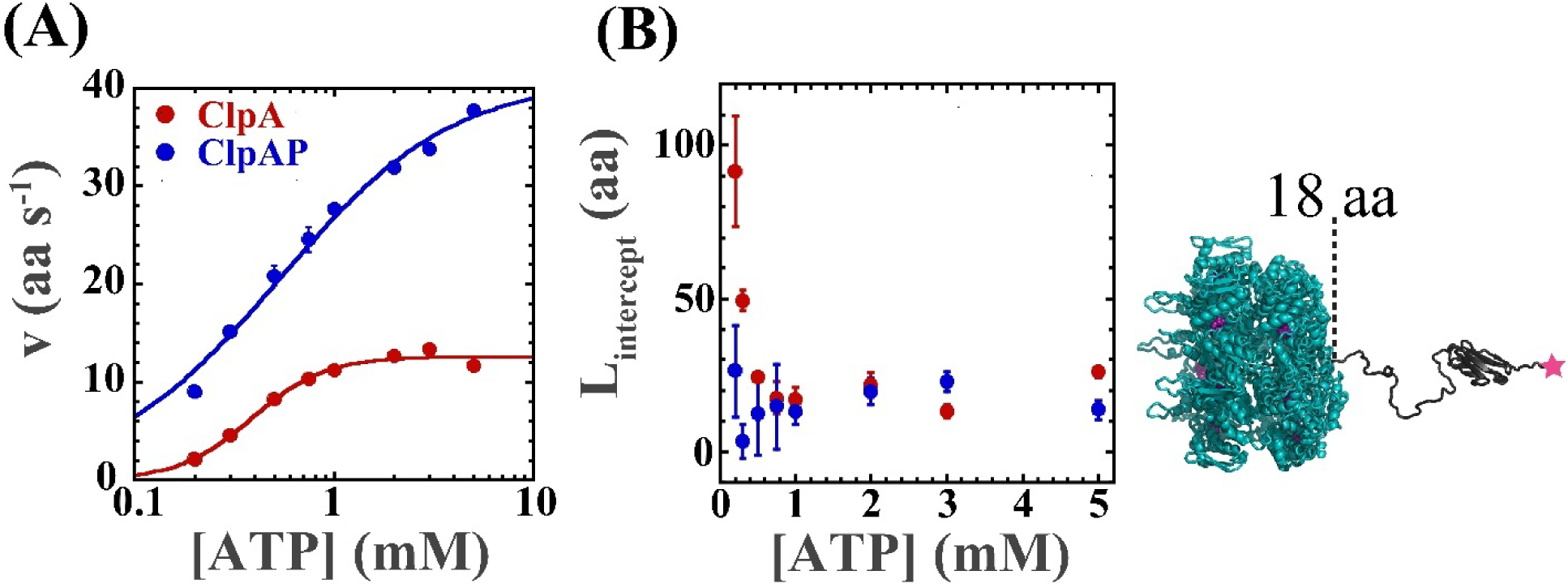
Results from peak time analysis. **(A)** The velocities, *v*, are [ATP]-dependent. Error bars in plots: S.E. The solid lines are the results of NLLS fits to the cooperative model for ClpA (red) and the noncooperative model for ClpAP (blue). Fitting yields *v_max_* = (12.6 ± 0.4) aa s^-1^, *K_d_* = (0.39 ± 0.02) mM, and *h* = (2.4 ± 0.3) for ClpA. For ClpAP, *v_max_* = (41.0 ± 1.2) aa s^-1^, and *K_d_* = (0.53 ± 0.05) mM. These values represent (parameter ± S.E.). **(B)** (*L_intercept_* ± S.E.) vs. [ATP]. All values are within error except for ClpA at 200 and 300 µM ATP, which are outliers. Average *L_intercept_* = (18 ± 5) aa, where the error is S.D. We interpret *L_intercept_* to be the length excluded from ATP-driven translocation, as illustrated on the right, where ClpA starts translocating with its front at 18 aa.

For ClpA, *v* depends cooperatively on [ATP] with a Hill coefficient, *h* ≈ 2.4, a maximum velocity at saturating [ATP], *v_max_*≈ 13 aa s^-1^, and an ATP-dissociation equilibrium constant, *K_d_* ≈ 0.39 mM, see Materials and Methods, Figure S3 (A), and Table S1 for nonlinear least-squares (NLLS) analysis. ∼13 aa s^-1^ on folded substrates is slower than ∼20 aa s^-1^ reported for unstructured substrates [12], suggesting that ClpA-catalyzed translocation at saturating [ATP] is slowed by folded domains.

In contrast, for ClpAP, *v* does not depend cooperatively on [ATP]. For ClpAP, we determined *v_max_* ≈ 41 aa s^-1^, and *K_d_* ≈ 0.53 mM, see Materials and Methods, Figure S3 (B), and Table S1. Comparing *v_max_* for folded substrates with ∼36 aa s^-1^ reported for unstructured substrates [13], we conclude that the Titin I27 domains do not impede translocation by ClpAP at saturating [ATP]. The observation that the velocity of ClpA-catalyzed unfolding and translocation depends cooperatively on [ATP], and the velocity of ClpAP-catalyzed unfolding and translocation does not, is identical to what was observed with unstructured substrates [12, 39].

The average *L_intercept_* for ClpA and ClpAP from the peak time analysis is (18 ± 5) aa, see Figure 4 (B). We have interpreted *L_intercept_* as the excluded length due to substrate binding. ∼18 aa seems reasonable for the average excluded length, given that a minimum of 15 aa is required for ClpA or ClpAP to initiate substrate unfolding and translocation on RepA_1-70_ [21].

At 200 and 300 µM ATP, *L_intercept_* for ClpA systematically increases to ∼49 and 91 aa respectively, so these values were excluded when calculating the average *L_intercept_*. It seems unlikely that the increase in *L_intercept_*would be due to an increase in excluded length since the assembly protocol was identical at all [ATP]. We hypothesize that at low [ATP], unfolding is much slower than translocation, artificially increasing *L_intercept_* from the excluded length.

### n-Step Sequential Mechanisms Describe Time Courses

The lag-phase in a single-turnover time course indicates at least two repeated steps with similar rate constants [12, 31, 32]. We interpret the peak and decay in the time courses to indicate arrival at the dye and induction of PIFE, followed by slow dissociation and loss of PIFE.

To model these phenomena, we propose Scheme 1 in Figure 5 (A). Here, ClpA starts pre-bound to RepA-Titin_X_. Upon mixing with ATP, ClpA takes a step with a rate constant, *k_obs_,* to form the first intermediate, *I_1_*. Then ClpA cycles through multiple steps with the same rate constant, *k_obs_*, to form the second, third, and eventually, *n*^th^ intermediate. Slow dissociation with a rate constant, *k_end_*, follows.

**Figure 5:**
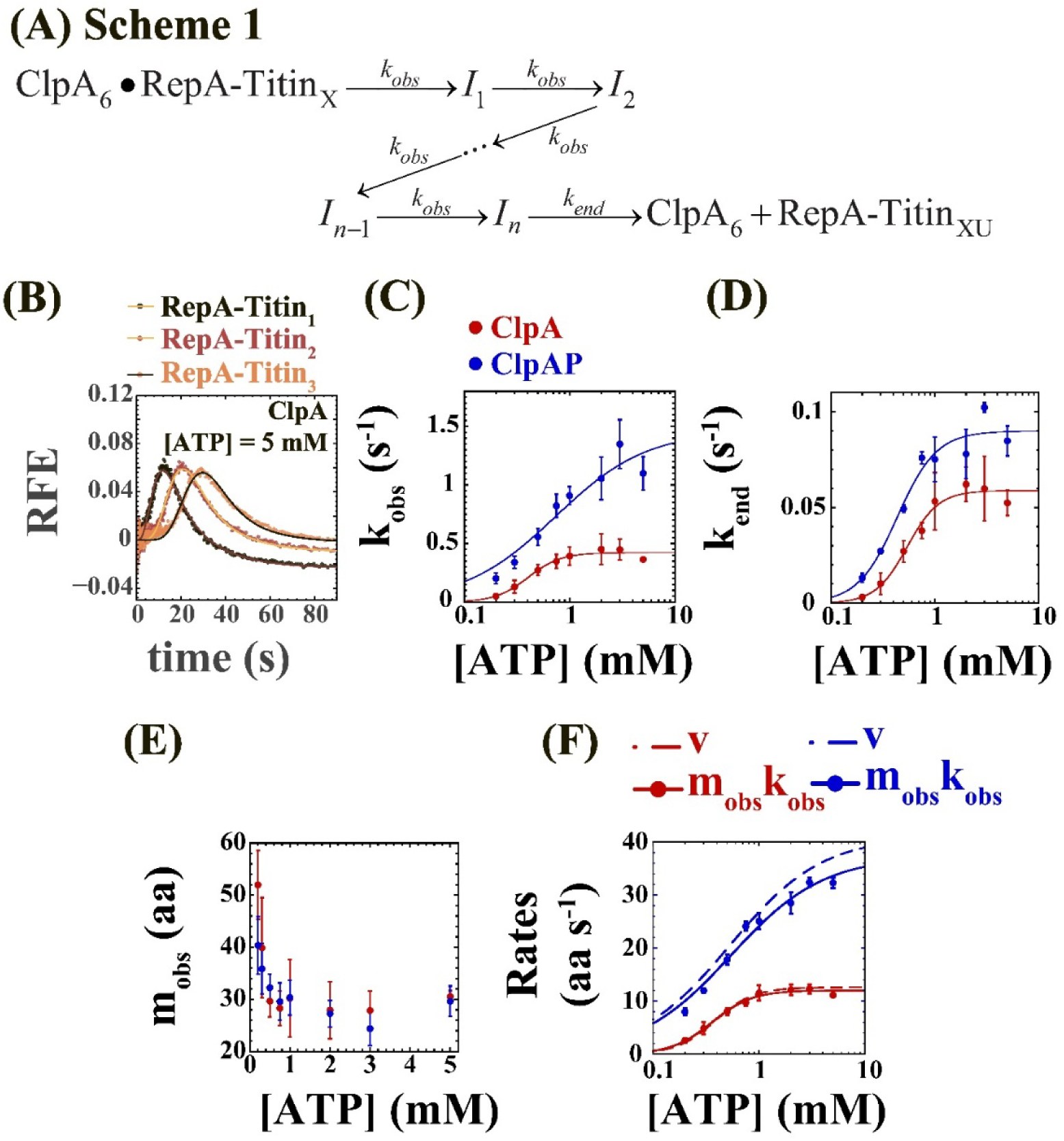
Mechanistic differences in ClpA- and ClpAP-catalyzed unfolding and translocation. **(A)** Scheme 1 is the simplest scheme describing all time courses. **(B)** Representative time courses for ClpA at 5 mM ATP (dots) fit to Scheme 1 (Solid lines). **(C)** *k_obs_* vs. [ATP] for ClpA (red dots) fit to the cooperative model (red line) with (*k_obs_)_max_* = (0.42 ± 0.02) s^-1^, *K_d_* = (0.40 ± 0.04) mM, and *h* = (2.8 ± 0.6). *k_obs_* vs. [ATP] for ClpAP (blue dots) fit to the noncooperative model (blue line) with (*k_obs_)_max_* = (1.46 ± 0.15) s^-1^, and *K_d_* = (0.74 ± 0.22) mM. **(D)** *k_end_* vs. [ATP] for ClpA (red dots) fit to the cooperative model (red line) with (*k_end_)_max_* = (0.059 ± 0.003) s^-1^, *K_d_* = (0.54 ± 0.05) mM, and *h* = (2.9 ± 0.6). *k_end_* vs. [ATP] for ClpAP (blue dots) fit to the cooperative model (blue line) with (*k_end_)_max_* = (0.090 ± 0.006) s^-1^, *K_d_* = (0.43 ± 0.05) mM, and *h* = (2.3 ± 0.6). **(E)** The kinetic step-size, *m_obs_*, increases from ∼29 aa at high [ATP] to ∼52 aa for ClpA and to ∼40 aa for ClpAP at low [ATP]. **(F)** Overall rates, *m_obs_k_obs_* vs. [ATP] for ClpA (red dots) fit to the cooperative model (solid red line) with (*m_obs_k_obs_)_max_* = (12.0 ± 0.4) aa s^-1^, *K_d_* = (0.36 ± 0.02) mM, and *h* = (2.3 ± 0.3). *m_obs_k_obs_* vs. [ATP] for ClpAP (blue dots) fit to the noncooperative model (solid blue line) with (*m_obs_k_obs_)_max_* = (40.0 ± 1.1) aa s^-1^, and *K_d_* = (0.59 ± 0.05) mM. Dashed lines show fitted curves for *v* vs. [ATP] from Figure 4 (C) for ClpA (red) and ClpAP (blue). **(C)-(F)** Error bars in plots: S.D. over three replicates. Error in parameters associated with cooperative and noncooperative models: S.E.

PIFE is expected when ClpA comes within ∼3 nm of AF555 [34–36], so it will occur at *I_n_* and, potentially, some number of intermediates before *I_n_*. We decided on a fitting function embodying Scheme 1 and including contributions from *I_n-1_*, *I_n_*, and RepA-Titin_XU_ based on analyses presented in Figure S4-S10. Time courses for all substrates at each [ATP] were globally fit using this fitting function and the assumption that, 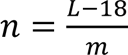. Here, *m* is the kinetic step-size, which is the observed number of aa translocated between two rate-limiting steps. (*L*-18) accounts for the excluded length due to pre-binding the substrate.

Figure 5 (B) shows a representative global fit of time courses from all three substrate lengths collected using ClpA at 5 mM ATP. The same global fits were performed for both ClpA and ClpAP at each [ATP]. The resulting fit parameters *k_obs_*, *k_end_*, and *m_obs_*, see Table S2, were averaged over three replicates.

The observed repeating rate constant, *k_obs_*, is [ATP]-dependent for both ClpA and ClpAP, see Figure 5 (C). Like the velocities determined in the peak time analysis, *k_obs_* depends cooperatively on [ATP] for ClpA but not ClpAP, see Figure S11 (A), (D), and Table S3. At all [ATP], *k_obs_* is faster for ClpAP compared to ClpA.

*k_end_* vs. [ATP] follows the cooperative model for both ClpA and ClpAP, indicating that substrate release is coupled to ATP binding, see Figure 5 (D), Figure S11 (B), (E), and Table S3.

For both ClpA and ClpAP, the kinetic step-size, *m_obs_*, is constant at *∼*29 aa, above 300 μM ATP. Below 300 μM ATP, *m_obs_* increases, see Figure 5 (E). This is different from our previously observed [ATP]-independent kinetic step-sizes of *∼*14 aa and *∼*5 aa for ClpA- and ClpAP-catalyzed translocation of unstructured substrates, respectively.

For ClpA, the overall rates, *m_obs_k_obs_*, are within error of the estimated velocities, *v*, from *L* vs. *t_peak_* plots, see Figure 5 (F) and 4 (A), indicating good agreement between model-independent analysis and global fitting. For ClpAP, the values are close but slightly deviate.

### Unfolding and Translocation are Differentially Coupled to ATP Binding

Scheme 1 shows a single repeated rate-limiting step of unknown identity. The [ATP]-dependence of the observed rate constant, *k_obs_*, reveals that this step is coupled to ATP binding, see Figure 5 (C).

In previous studies on unstructured substrates, the kinetic step-size represented translocation in the absence of unfolding [12, 13]. Here, the kinetic step-sizes observed on folded substrates are larger than step-sizes for translocation alone, see Figure 5 (E). The process observed here is likely more complicated than Scheme 1. Rather, Scheme 2, shown in Figure 6 (A), reflects the repeated unfolding and translocation processes catalyzed by ClpA or ClpAP on a folded substrate. Here, *k_U_* is the rate constant of unfolding, and *k_T_* is the rate constant of translocation. Each unfolding event with a step-size, *m_U_*, is followed by multiple translocation steps with a smaller step-size, *m_T_*, until the next folded domain is reached. The single-turnover time courses are sensitive to the slowest repeating step, raising the question-are the time courses reporting on translocation, unfolding, or a combination of both?

**Figure 6:**
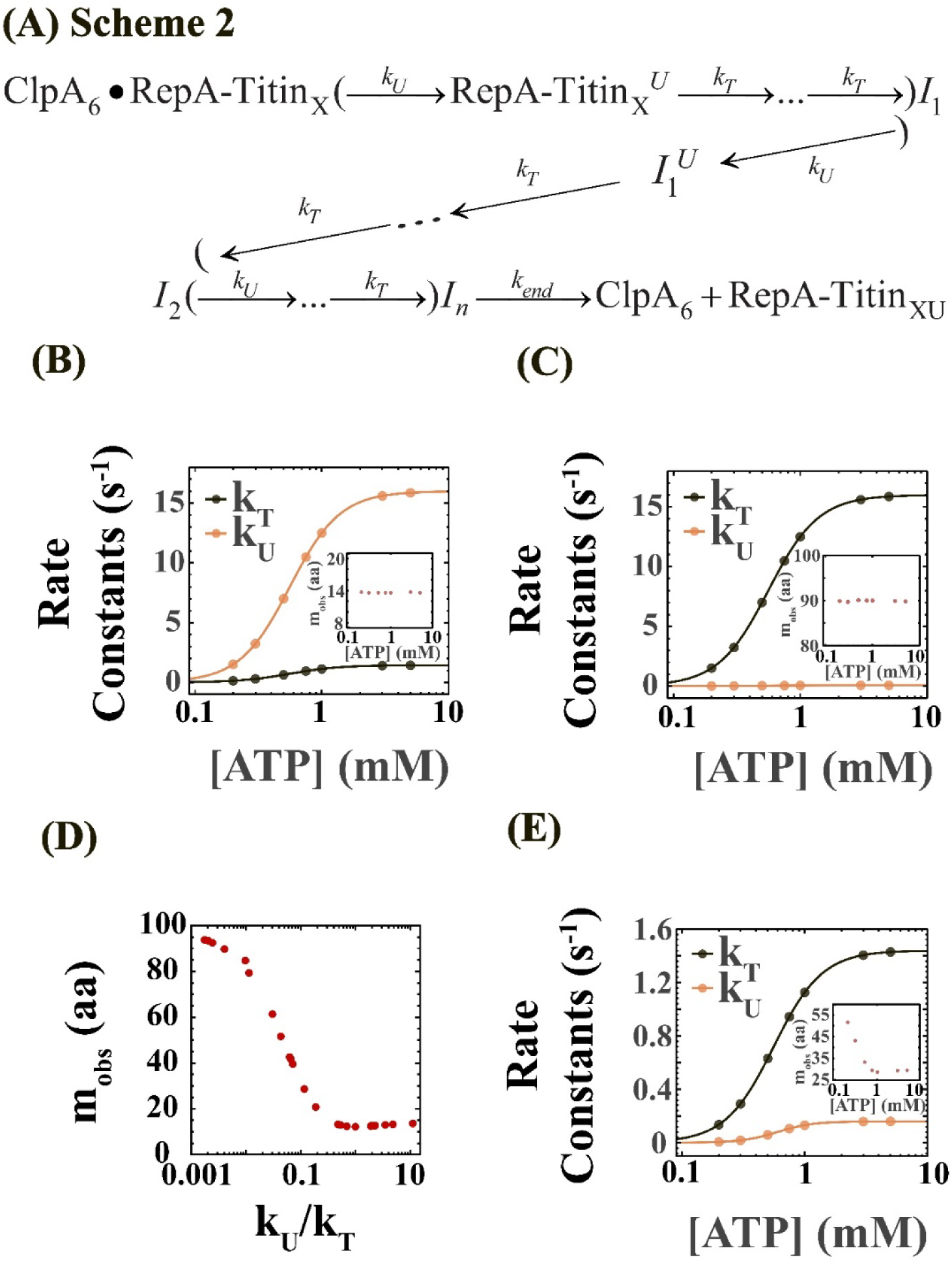
The way unfolding and translocation are kinetically coupled to ATP binding predicts patterns in the observed kinetic step size, *m_obs_*. (A) Scheme 2 describes the underlying mechanism of ClpA-or ClpAP-catalyzed protein unfolding and translocation with the translocation rate constant, *k_T_*, and the unfolding rate constant, *k_U_*. **(B)** We simulated time courses with Scheme 2, using *m_T_* = 14 aa, *m_U_* = 97 aa, *k_T_* from the black curve, and *k_U_* from the yellow curve. Here, translocation is rate-limiting (*k_U_* >> *k_T_*) at all [ATP]. Fitting the simulated time courses via Scheme 1 predicts, *m_obs_* ≈ *m_T_*, constant with [ATP], see inset. **(C)** The same analysis as (B) with unfolding being rate-limiting (*k_T_* >> *k_U_*) at all [ATP] predicts *m_obs_* ≈ *m_U_*, also constant with [ATP], see inset. (D) *m_obs_* varies with *k_U_*/*k_T_*. **(E)** The same analysis as (B)-(C) with *k_T_* > *k_U_* at all [ATP] and *k_U_*/*k_T_* decreasing at low [ATP], predicts *m_obs_* to increase at low [ATP], see inset.

If unfolding is rate-limiting, we should detect the unfolding step-size, *m_U_*. If translocation is rate-limiting, we should detect the translocation step-size, *m_T_*. If one could isolate each process, then *k_U_*and *k_T_* would separately exhibit [ATP]-dependencies.

To identify the process observed at different [ATP], we used Scheme 2 to simulate time courses using hypothetical [ATP]-dependencies for *k_U_*, and the previously published [ATP]-dependencies for *k_T_*, see Figure 6 (B). Here, translocation is rate-limiting (*k_U_* >> *k_T_*) at all [ATP]. Because Titin I27 cooperatively unfolds in a single step [17, 23, 40], we set the unfolding step-size, *m_U_* = 97 aa, which represents the length of a single Titin I27 domain, see Figure 1 (A). The translocation step-size, *m_T_*, was set to values previously reported by us for translocation on unstructured substrates, i.e., ∼14 and ∼5 aa for ClpA and ClpAP, respectively [12, 13]. Fitting simulated time courses using Scheme 1 yields the predicted kinetic step-size, *m_obs_*, see inset in Figure 6 (B). We repeated this analysis in Figure 6 (C) so that unfolding is rate-limiting (*k_T_* >> *k_U_*) at all [ATP].

When either unfolding or translocation is rate-limiting at all [ATP], the same rate-limiting step should be detected at all [ATP], resulting in an [ATP]-independent kinetic step-size [31, 41]. Consistently, *m_obs_* vs. [ATP] is [ATP]-independent in Figure 6 (B)-(C), with *m_obs_* ≈ *m_T_* when translocation is rate-limiting, and *m_obs_* ≈ *m_U_* when unfolding is rate-limiting, see insets.

Experimentally, we observe an increase in step-size at low [ATP], see Figure 5 (E), which rules out cases like Figure 6 (B)-(C). Instead, an [ATP]-dependent kinetic step-size implies a shift in the observed rate-limiting step with changing [ATP] [31, 41]. Thus, an [ATP]-dependent kinetic step-size predicts that the isotherms describing [ATP]-dependencies for *k_U_* and *k_T_* either cross or approach with changing [ATP]. Another way to think about this is that the *k_U_*/*k_T_* ratio is changing as a function of [ATP].

To test this, we simulated time courses by varying *k_U_*/*k_T_* in Scheme 2, subjected them to global fitting using Scheme 1, and determined the predicted step-size, *m_obs_*. We found that, as *k_U_*/*k_T_* decreases, *m_obs_* increases from a value close to the smaller translocation step-size, *m_T_*, to the larger unfolding step-size, *m_U_*, see Figure 6 (D) and Figure S12. Time courses simulated with *m_T_* = 14 aa and *m_U_* = 97 aa yield *m_obs_* = 29 aa when *k_U_*/*k_T_* = 0.11, see Figure 6 (D). Time courses simulated with *m_T_* = 5 aa and *m_U_* = 97 aa yield *m_obs_* = 29 aa when *k_U_*/*k_T_* = 0.04, see Figure S12. Experimentally, we detect *m_obs_* ≈ 29 aa for both ClpA and ClpAP at high [ATP], see Figure 5 (E).

Figure 6 (D) shows that when *m_T_* = 14 aa, *m_U_* = 97 aa, and *k_U_*/*k_T_* = 0.04, we get *m_obs_* ≈ 52 aa, which is the kinetic step-size detected for ClpA at low [ATP], see Figure 5 (E). Figure S12 shows that when *m_T_* = 5 aa, *m_U_* = 97 aa, and *k_U_*/*k_T_* = 0.02, we get *m_obs_* ≈ 40 aa, which is the kinetic step-size detected for ClpAP at low [ATP], see Figure 5 (E).

To test if we can predict the trends for ClpA in Figure 5 (E) using the *k_U_*/*k_T_* ratio, we generated the theoretical curve for *k_U_* in Figure 6 (E), where *k_U_* = 0.11*k_T_* at high [ATP] and *k_U_*= 0.04*k_T_* at low [ATP]. Next, we generated simulated time courses at each [ATP] with these *k_U_* values, *m_T_* = 14 aa, *m_U_* = 97 aa, and *k_T_* from the experimentally determined isotherm for ClpA-catalyzed translocation [12]. From the simulated time courses, we determined *m_obs_*, see inset of Figure 6 (E). The predicted *m_obs_* vs. [ATP] closely matches the trend observed experimentally in Figure 5 (E). A similar analysis predicts the trend in *m_obs_* vs. [ATP] seen for ClpAP in Figure 5 (E) by assuming *k_U_*= 0.04*k_T_* at high [ATP] and *k_U_* = 0.02*k_T_*at low [ATP].

In summary, we propose the model in Figure 7 to describe the experimental observations. First, ClpA binds the folded substrate, RepA-Titin_3_, with the first 18 aa of RepA_1-70_ occluded in the axial channel. Upon binding ATP, translocation occurs with a rate constant, *k_T_*, until ClpA encounters a folded domain at 74 aa. There, unfolding is catalyzed with a rate constant, *k_U_* < *k_T_*. Unfolding is followed by several translocation steps until the next fold is encountered at 171 aa. This slow-unfolding-fast-translocation cycle repeats until ClpA reaches AF555 at 362 aa, inducing PIFE. Slow dissociation of ClpA from the unfolded substrate ensues, which also requires ATP binding/hydrolysis. Although Figure 7 is drawn with ClpA, the same model applies to ClpAP.

**Figure 7:**
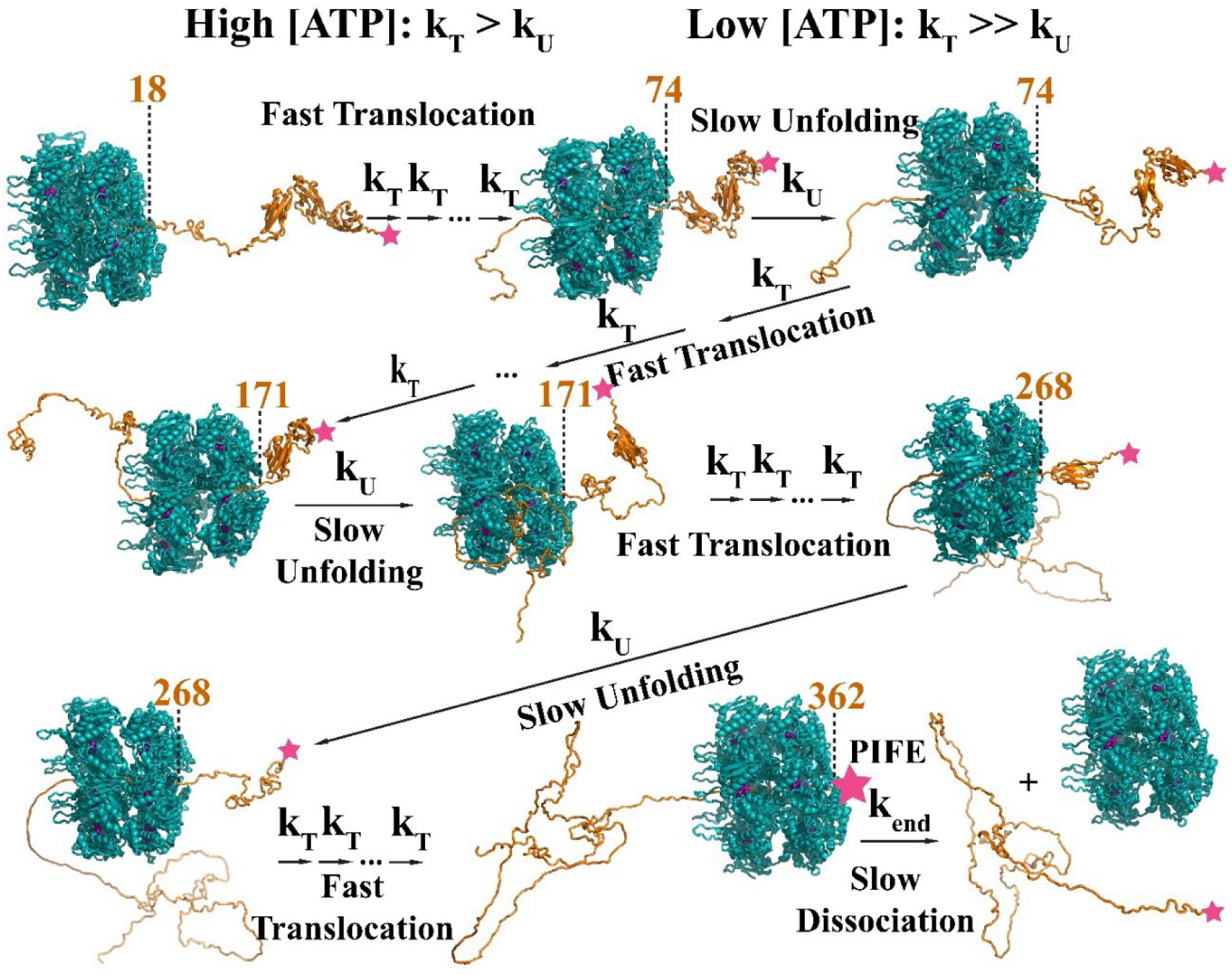
Model for ClpA unfolding and translocating on a folded substrate. Numbers indicate the position of the ClpA front along the substrate primary sequence. ClpA (teal) binds the unstructured RepA_1-70_ sequence. Upon ATP addition, ClpA unfolds 97aa of RepA-Titin_3_ at the rate constant, *k_U_*, followed by multiple translocation steps at the rate constant, *k_T_*. The process repeats. Both *k_U_* and *k_T_* decrease with [ATP]. Partial rate-limiting kinetics yields *m_obs_* ≠ *m_U_* or *m_T_*, but something in between. After complete unfolding and translocation of all Titin I27 domains, ClpA reaches AF555 (pink star) to induce PIFE, then dissociates slowly at the rate constant, *k_end_*. We propose a similar mechanism for ClpAP, with the motor degrading the unfolded chain as it advances.

## Discussion

### Substrate Folds Impede ClpA-Catalyzed Translocation

We originally developed the single-turnover fluorescence stopped-flow method to determine the mechanism of ClpA-catalyzed polypeptide translocation. Initially, we used unstructured substrates, ensuring that the time courses reported only on translocation. We found that, for ClpA at saturating [ATP], the translocation rate constant is ∼1.4 s^-1^, and the kinetic step-size is ∼14 aa, yielding an overall rate of ∼19 aa s^-1^ [12]. Using the same method, we found that for ClpAP at saturating [ATP], the translocation rate constant is ∼7.9 s^-1^, and the kinetic step-size is ∼5 aa, yielding an overall rate of 36 aa s^-1^ [13].

Unlike ClpA, ClpAP proteolytically degrades the translocated substrate. However, our method is sensitive to interactions of the motor with the fluorophore and does not directly report on degradation. Differences in the kinetic parameters for ClpA and ClpAP revealed that ClpP allosterically modulates ClpA activity [13].

Here we show that substrate fold significantly impacts translocation. At saturating [ATP], ClpA or ClpAP translocates and unfolds with observed rate constants of ∼0.4 s^-1^ or ∼1.5 s^-1^, respectively. These values are lower than values reported for ClpA or ClpAP-catalyzed translocation of 1.4 s^-1^ or 7.9 s^-1^, respectively, see Table S4. Also, for ClpA, the overall rate decreases from ∼20 aa s^-1^ on unstructured substrates [12] to ∼12 aa s^-1^ on folded substrates. So, the motor does not simply translocate unimpeded by the substrate fold. In contrast, ClpAP exhibits comparable overall rates of ∼36 aa s^-1^ [13] and ∼40 aa s^-1^ on unstructured and folded substrates, respectively. This could indicate that the overall process in ClpAP-catalyzed unfolding and translocation is limited by translocation rates, translocation being unimpeded by substrate fold. This raises several questions regarding how and why the presence of the substrate affects the ability of the motor component to unfold. Nevertheless, this observation is consistent with previous reports on the rates of ClpAP-catalyzed degradation being equal on folded and unstructured substrates [9].

ClpA is slower than ClpAP in both unfolding and translocation [13, 37, 42, 43]. Moreover, on both folded and unstructured substrates translocated by ClpA, the repeating rate constant exhibits a cooperative dependence on [ATP]. This cooperativity indicates that at least two ATP binding and hydrolysis sites are cooperating to couple ATP binding/hydrolysis to translocation and unfolding. In contrast, ClpAP shows no such cooperativity, indicating that fewer ATP binding/hydrolysis sites are needed to catalyze the unfolding and translocation reactions. So, the proteolytic component, ClpP, allosterically influences the way ClpA coordinates and couples ATP binding and hydrolysis to repeating rounds of unfolding and translocation.

### Coordinated Functions of the Two ATP Binding Sites

ClpA has two Nucleotide Binding Domains (NBDs). Upon binding ATP, or ATP analogs in these NBDs, ClpA assembles into double-ring hexamers. The rings are termed domain 1 (D1) and domain 2 (D2), indicating that Nucleotide Binding Domain 1 (NBD1) resides in the D1 ring and Nucleotide Binding Domain 2 (NBD2) resides in the D2 ring [6, 10, 44, 45]. ATP binding at NBD1 is crucial for hexamerization [44, 46, 47], while NBD2 catalyzes the most ATP hydrolysis. D1-active-D2-inactive ClpA (ClpA^D2^) associated with ClpP can degrade less stable substrates. More stable substrates need activity in both rings [48].

In cryo-EM structures of ClpAP, both D1 and D2 engage the substrate at 2 aa intervals in the axial channel through pore loops extending from each NBD [29, 38, 45, 49, 50]. The dominant structural model is that ATP binding and hydrolysis induce these pore loops to toggle between up and down conformations, translocating the substrate through the channel [51, 52]. From this model, the step-size of translocation was proposed to be 2 aa [29, 51]. However, we and others have yet to detect a step-size in solution as small as 2 aa step^-1^ [12, 13, 53].

Our previous estimates on kinetic step-sizes led us to propose a model where the two NBDs coordinate with different mechanisms during ClpA-vs. ClpAP-catalyzed translocation [13, 54]. Both NBDs start in the ATP-bound state, engaging the substrate via pore loops. ATP hydrolysis first occurs in NBD1, translocating the substrate further into the axial channel. Then the substrate affinity of the associated pore loop decreases [52], allowing it to reengage the substrate at a new position. Finally, a new ATP binds at NBD1, rebooting the cycle.

Based on several observations, we proposed that the step-size of this process catalyzed at D1 repeats every 14 aa translocated [12, 39, 55, 56]. Next, D2 catalyzes ATP hydrolysis and translocates the substrate in 5 aa steps, translocating the substrate out of the axial channel [13, 55]. Because the step-size of D1-catalyzed translocation is larger than that of D2, we proposed that the substrate will first pucker in the axial channel during translocation by D1. Subsequently, multiple translocation steps need to be catalyzed by D2 before D1 takes another step [13, 54]. Consistently, the events catalyzed at D1 are slower than at D2. Consequently, multiple 5 aa steps take place at D2 before a single 14 amino acid step at D1 occurs. With RepA-Titin_X_ substrates in hand, we are now well-positioned to test this model using ATPase-activity-deficient mutants of ClpA.

Here we show that ClpA and ClpAP-catalyzed unfolding and translocation have comparable kinetic step-sizes defined by the *k_U_*/*k_T_*ratio. Lowering [ATP] reduces both *k_U_* and *k_T_*, with a greater effect on *k_U_*. Since both D1 and D2 are necessary for unfolding stably folded substrates [48], *k_U_* should decrease in ClpA Walker B mutants [57–59] deficient in ATP hydrolysis but not ATP binding at each NBD. Since D2 is most important in ATP hydrolysis and unfolding, we expect ClpA^D2^ to unfold and translocate more slowly compared to D1-inactive-D2-active ClpA (ClpA^D1^). Both processes should be differentially coupled to ATP binding for these mutants. We hypothesize that *k_U_*/*k_T_*would decrease in both mutants, resulting in increased kinetic step-sizes compared to wildtype.

Understanding how unfolding and translocation are coupled to ATP binding in ClpA and ClpA will also allow us to test whether the cooperativity observed in ClpA is inter- or intra-monomeric. Cryo-EM structures point to ATP hydrolysis in adjacent monomers [29], which we predict to be associated with inter-monomeric cooperativity.

### Unfolding as the Slower Process

Single-molecule optical trapping experiments on ClpAP-catalyzed unfolding and translocation of wild-type Titin I27 domains revealed that at 5 mM ATP, the time constant of unfolding is ∼0.8 s. Also, one unfolding step is followed by ∼20 translocation steps with an average time constant of ∼0.55 s in the N-to-C direction. Thus, the average time spent translocating an entire unfolded domain should be ∼20 × 0.55 s ≈ 11 s, leading to the conclusion that slow translocation follows rapid unfolding [17].

At the same [ATP], our stopped-flow experiments yielded a kinetic step-size of ∼29 aa for ClpAP on folded substrates. As shown in the Results, if the unfolding step size, *m_U_* ≈ 97 aa, [17, 23, 40], and the translocation step-size, *m_T_*≈ 5 aa [13], then a ∼29 aa step size corresponds to *k_U_*/*k_T_* ≈ 0.04. On unstructured substrates, *k_T_* ≈ 7.6 s^-1^ [13], so *k_U_* ≈ 0.04*k_T_* ≈ 0.3 s^-1^. So, the average time spent per translocation step, 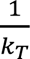 ≈ 0.13 s, and per unfolding step, 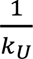 ≈ 3.3 s. Given that a single Titin I27 domain is 97 aa, there are 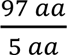 ≈ 19 translocation steps per unfolding step. Thus, based on our measured parameters, we estimate the average time ClpAP spends translocating an unfolded Titin I27 domain in the stopped-flow assay to be 19 × 0.13 ≈ ≈ 2.5 s. Thus, on average, ClpAP spends ∼3.3 s unfolding a Titin I27 domain, followed by ∼2.5 s to translocate the newly unfolded chain completely. Our observations are consistent with unfolding and translocation being comparable in rate.

The differences between optical trapping and our single-turnover stopped-flow measurements may have multiple sources. First, on RepA-Titin_X_ substrates, we do not uniquely detect either translocation or unfolding. We are assuming that translocation on a newly unfolded Titin I27 domain would follow the same mechanism as on our previously reported unfolded substrates, even though the directionality is different. This may be a good assumption since it has been reported that ClpAP-catalyzed unfolding and translocation on a freshly unfolded Titin I27 domain is only modestly slower N-to-C compared to C-to-N [17].

The differences in experimental conditions likely have the biggest impact on the rates. The optical trapping experiments were carried out at 18-20 °C [17] compared to 25 °C used here [13]. Also, the applied force in optical trapping impacts the energy landscape and the rate constants [60].

### Motor-Specific Differential Coupling

The RepA-Titin_X_ substrates used here were previously used to examine substrate unfolding and translocation catalyzed by the ClpB disaggregase [22, 23]. ClpA and ClpB are AAA+ members sharing 42 % sequence identity and 64 % sequence similarity [61]. Both are responsible for viability and virulence in bacteria. Unlike ClpA, ClpB does not associate with any protease [62].

Using the same substrates and methods, we can, for the first time, compare the mechanisms of ClpA- and ClpB-catalyzed protein unfolding. We have reported that the kinetic step-size of ClpB-catalyzed protein unfolding is ∼97 aa at saturating [ATP][23]. We interpreted this to indicate that unfolding was completely rate-limiting and that ClpB catalyzes cooperative unfolding of a Titin I27 domain, followed by fast and undetectable translocation [17, 23, 40].

For ClpB at low [ATP], the kinetic step-size drops below 97 aa, suggesting that unfolding and translocation have become partially rate-limiting [23]. In contrast, unfolding and translocation are partially rate-limiting for ClpA at saturating [ATP]. For ClpA, the observed kinetic step-size for unfolding and translocation increases at low [ATP] as unfolding becomes more rate-limiting. Taken together, our results point to motor-specific differential coupling of unfolding and translocation to ATP binding.

## Acknowledgments

We thank Zachary G. Cuny, Wade Bexley, Dr. Nate W. Scull, and Dr. Kaila B. Fuller for purifying ClpA; Dr. Elizabeth C. Duran and Dr. Clarissa L. Durie for purifying ClpP; and Ryan M. Requijo for purifying RepA-Titin_3_. This work was supported by the National Institute of General Medical Sciences of the National Institutes of Health under award no. R35GM156375 (to A.L.L). Computational work was performed using the University of Alabama at Birmingham Supercomputer Cheaha, which is supported in part by the National Science Foundation under grant no. OAC-1541310, the University of Alabama at Birmingham, and the Alabama Innovation Fund.

## Author Contributions

Conceptualization, L.I. and A.L.L.; resources, L.I., J.K.B. and A.L.L.; data curation, L.I.; formal analysis, L.I. and A.L.L; methodology, L.I., and A.L.L.; investigation, L.I. and A.L.L.; validation, L.I. and A.L.L.; writing – original draft, L.I.; writing – review & editing, A.L.L.; supervision, A.L.L.

