## Supplemental Information for "ClpA- and ClpAP-Catalyzed Unfolding and Translocation are Differentially Coupled to ATP Binding"

### S. Supplemental Section

#### S.1. Peak Time Analysis for ClpAP

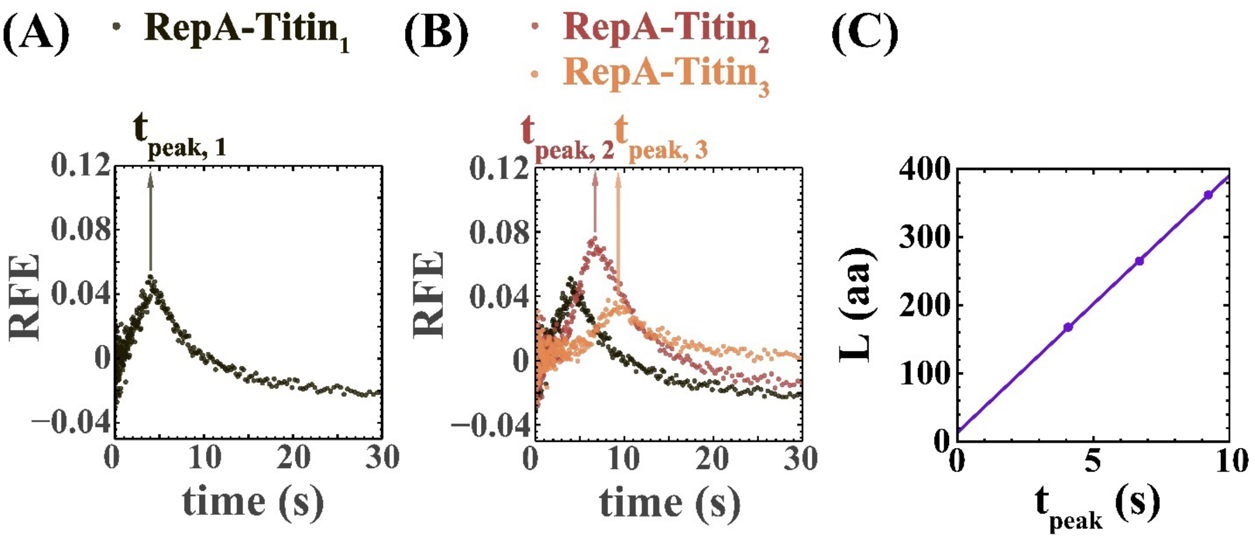


**Figure S1:** **(A)** Peak time analysis for ClpAP at [ATP] = 5 mM. **(A)** Representative time course from RepA-$Titin_1$ displaying a lag, a peak, and a decay. **(B)** Time courses from all RepA-$Titin_X$ substrates at the same [ATP]. **(C)** Substrate length, $L$, vs. average peak times, $t_{peak}$, from three replicates. The vanishingly small error bars represent standard deviations (S.D.) in $t_{peak}$. The straight line represents a linear-least-squares fit with a slope (velocity), $v$ = (37.7 ± 0.4) aa $s^{-1}$ and $L_{intercept}$ = (14 ± 3) aa, with the errors representing standard errors (S.E.).

#### S.2. Peak Time Analysis at Varying [ATP]

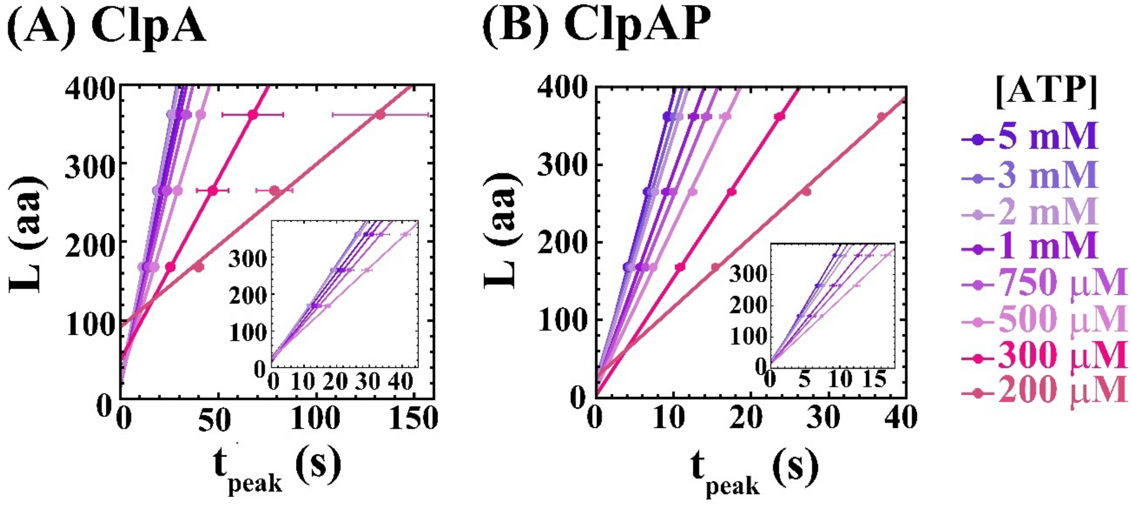


**Figure S2:** $L$, vs. $t_{peak}$ at all [ATP] for both **(A)** ClpA and **(B)** ClpAP, with the error bars representing S.D. of $t_{peak}$ for three replicates. The trends at each [ATP] are described by straight lines whose slopes (velocities) are plotted in Figure 4 (A), and intercepts are plotted in Figure 4 (B).

### S.3. $v$ vs. [ATP] Plots Reveal Cooperativity for ClpA but not ClpAP

For both ClpA and ClpAP, we have $n = 8$ data points per $v$ vs. [ATP] plot in Figure 4 (A). The number of parameters to be optimized when fitting the data with the noncooperative and cooperative models are $p_1 = 2$ and $p_2 = 3$, respectively, see Materials and Methods. The set of $p_1$ parameters is nested within the set of $p_2$ parameters. Figure S3 (A) and (B) show $v$ vs. [ATP] for ClpA and ClpAP, respectively, fit to both models.

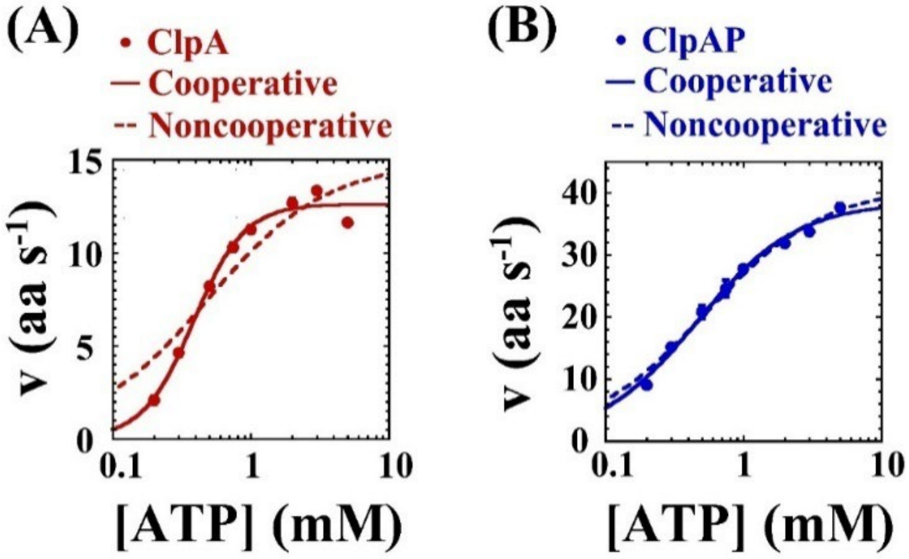


**Figure S3:** Plots of velocities, $v$, from Figure S2 and shown in Figure 4 (A) as a function of [ATP]. Both plots were fit to cooperative (solid lines) and noncooperative (dashed lines) models. All error bars represent S.E. of v. (A) For ClpA, the cooperative model is an improvement over the noncooperative model. (B) For ClpAP, the noncooperative model is sufficient to describe the trend.

Let the Residual Sum of Squares, $RSS$, when fitting $v$ vs. [ATP] be $RSS_1$ and $RSS_2$, for the noncooperative and cooperative models, respectively. Table S1 shows the fit parameters. Thus, we calculate the $F$-statistic, $F_{cal} = \frac{(\frac{RSS_1 - RSS_2}{p_2 - p_1})}{(\frac{RSS_2}{n - p_2})}$.

**Table S1:** Parameters obtained by fitting $v$ vs. [ATP] (from peak time analysis) to cooperative and noncooperative models. The error reported is S.E. Rows highlighted in green represent the simplest model describing the trend as determined using an $F$-test.

| **ClpA** | | | | | | |
|---|---|---|---|---|---|---|
| **Model** | **No. of fit parameters, *p*** | $R^2$ | **Fit *RSS*** | $v_{max}$ (aa $s^{-1}$) | $K_d$ (mM) | $h$ |
| Noncooperative | 2 | 0.873 | 14.40 | 14.9 ± 1.5 | 0.48 ± 0.16 | 1 (constrained) |
| Cooperative | 3 | 0.984 | 1.79 | 12.6 ± 0.4 | 0.39 ± 0.02 | 2.4 ± 0.3 |
| **ClpAP** | | | | | | |
| **Model** | **No. of fit parameters, *p*** | $R^2$ | **Fit *RSS*** | $v_{max}$ (aa $s^{-1}$) | $K_d$ (mM) | $h$ |
| Noncooperative | 2 | 0.987 | 8.64 | 41.0 ± 1.2 | 0.53 ± 0.05 | 1 (constrained) |
| Cooperative | 3 | 0.991 | 6.24 | 38.5 ± 1.6 | 0.46 ± 0.04 | 1.2 ± 0.1 |

If the null hypothesis is that the model with $p_2$ parameters does not significantly improve the fit over the model with $p_1$ parameters, $F_{cal}$ will follow the $F$-distribution, with ($p_2$-$p_1$, $n$-$p_2$) degrees of freedom [1]. We reject the null hypothesis if $F_{cal} > F_{critical}$, with a false-rejection probability of $\alpha = 0.01$. This is how we determined that $v$ vs. [ATP] follows the cooperative model for ClpA and the noncooperative model for ClpAP.

### S.4. Fitting Individual Time Courses with *n*-Step Sequential Mechanisms to Determine the Number of Intermediates Contributing to RFE

Scheme 1 shown in Figure 5 (A) was hypothesized to describe all time courses. To determine the number of intermediates before the end contributing to the RFE, we fit individual time courses to Scheme 1 with contributions from the last intermediate, $I_n$, and the unfolded substrate, RepA-Titin$_{XU}$ using the following equation.

$$RFE(t) = \mathcal{L}^{-1}\left(A_n \frac{k_{obs}{}^n}{(k_{end}+s)(k_{obs}+s)^n} + A_{xU}\frac{k_{end}k_{obs}{}^n}{s(k_{end}+s)(k_{obs}+s)^n}\right)$$

(S-I)

Here $\mathcal{L}^{-1}$ is the inverse Laplace transform operator, $A_n$ is the signal from $I_n$, $A_{xU}$ is the signal from RepA-Titin$_{xU}$, and $s$ is the Laplace variable. See Figure S4 (A) for a time course collected using ClpA and RepA-Titin$_1$ at 5 mM ATP, fit using Eq. (S-I). The parameters being optimized are $k_{obs}$, $k_{end}$, $A_n$ and $A_{XU}$.

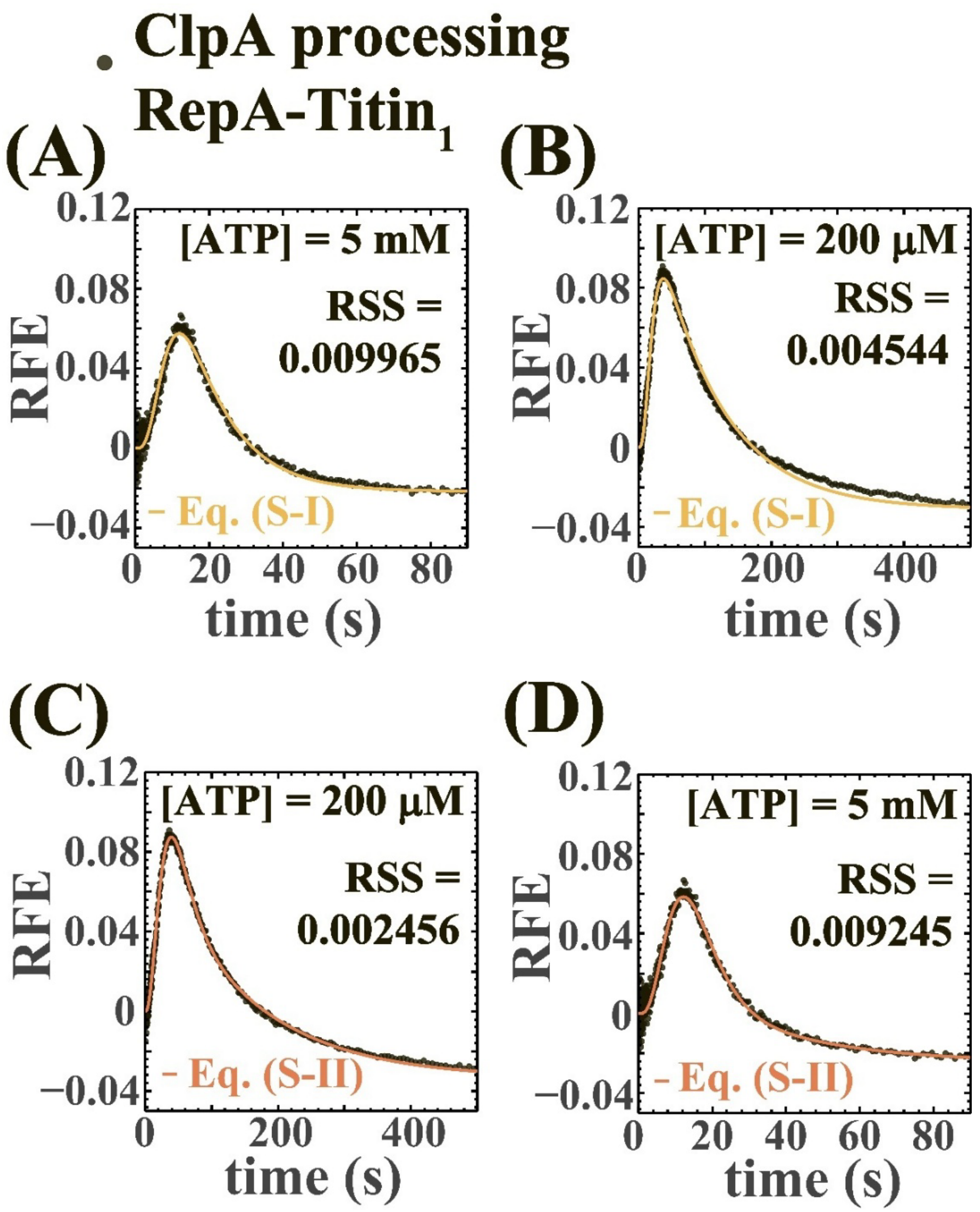


**Figure S4:** Time courses shown were collected using ClpA and RepA-Titin$_1$ at 5 mM, and 200 μM [ATP], the highest and lowest [ATP] tested, respectively. **(A)** Time course at high [ATP] fit to Eq. (S-I). **(B)** Fitting with Eq. (S-I) gives rise to systematic deviation in the decay phase at low [ATP]. **(C)** Eq. (S-II) reduces systematic deviation compared to (B). **(D)** Eq. (S-II) describes data at high [ATP] yet does not visually improve the fit compared to (A). Even then, an *F*-test involving the fit *RSS* values at a level of significance $\alpha = 0.01$ says that (D) and (C) are improvements over (A) and (B), respectively.

Eq. (S-I) describes the time course, so we proceeded to fit all time courses collected using ClpA with all substrates at all [ATP]. A time course collected using ClpA bound to RepA-Titin$_1$ at 200 μM [ATP] has been shown in Figure S4 (B), where the fitted curve undershoots the decay phase. This deviation in the decay phase is present at low [ATP] for other substrates as well (not shown). The deviation systematically increases as [ATP] decreases.

We hypothesized that contributions from one more intermediate, $I_{n-1}$, in the fitting function to get Eq. (S-II), may reduce the systematic deviations between time courses and fitted curves.

$$RFE(t) = \mathcal{L}^{-1}\left(A_{n-1}\frac{k_{obs}^{\ n-1}}{(k_{obs}+s)^n} + A_n\frac{k_{obs}^{\ n}}{(k_{end}+s)(k_{obs}+s)^n} + A_{xU}\frac{k_{end}k_{obs}^{\ n}}{s(k_{end}+s)(k_{obs}+s)^n}\right) \tag{S-II}$$

The extra parameter being optimized in this case, $A_{n-1}$, accounts for the signal from the penultimate intermediate, $I_{n-1}$. All other parameters being optimized are the same as in Eq. (S-I). Using Eq. (S-II), we observed a decrease in the systematic deviation between the decay phase and the trendlines at low [ATP], see Figure S4 (C). Consistent with the visual improvement of the fit, an $F$-test at a level of significance, $\alpha = 0.01$, indicates a statistically significant improvement when using Eq. (S-II) compared to Eq. (S-I). The $F$-test was carried out in a manner similar to Sup. Section S.1.

We proceeded to fit time courses collected at high [ATP] using Eq. (S-II). Even if there is no visual improvement in the fit shown in Figure S4 (D) compared to (A), an $F$-test with a level of significance, $\alpha = 0.01$, tells us that there is a statistically significant improvement in the fit. Taking this into account, and because it would be easier to compare phenomena at low and high [ATP] consistently if we fit time courses collected using all substrates at all [ATP] individually using the same model, we chose Eq. (S-II) for further analysis.

Similar analysis was carried out on data collected using ClpAP instead of ClpA to conclude that Eq. (S-II) describes the time courses best (fits not shown).

### S.5. Fitting Individual Time Courses to Identify Parameters that are Constant Across All Substrates

Individual time courses collected using ClpA bound to RepA-Titin$_X$, all collected at 5 mM [ATP], and fit to Eq. (S-II) are shown in Figure S5 (A)-(C). From this analysis, the repeating rate constant, $k_{obs}$, the terminal dissociation rate constant, $k_{end}$, and the number of steps, $n$, were plotted vs. substrate length, $L$, in Figure S5 (D)-(F).

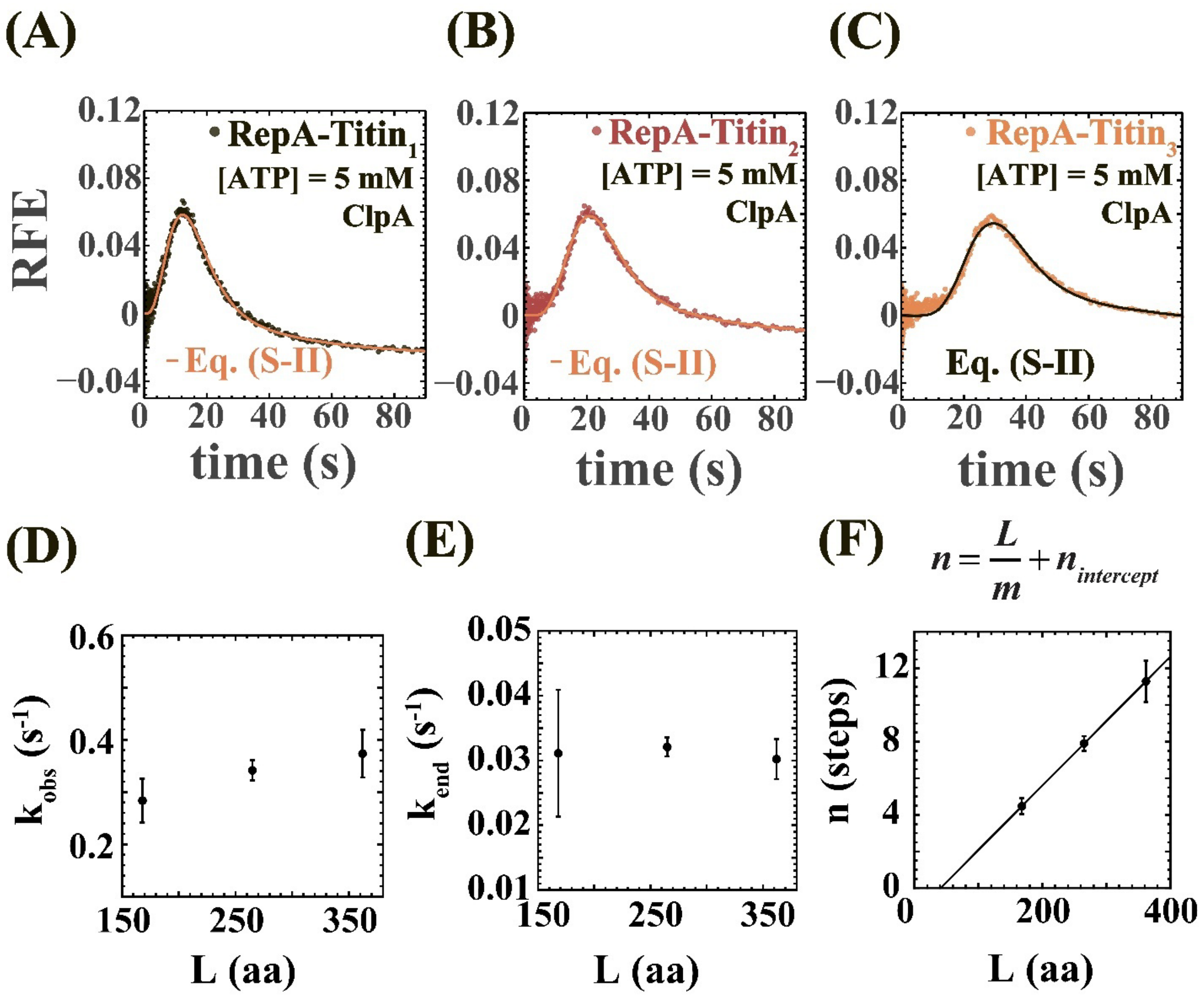


**Figure S5:** **(A)-(C)** Time courses from RepA-Titin1, RepA-Titin2 and RepA-Titin3, respectively, with ClpA at 5 mM ATP. All time courses were individually fit to Eq. (S-II). **(D)-(E)** $k_{obs}$ and $k_{end}$, respectively, of ClpA are within standard deviations for each substrate. **(F)** Number of steps, n, for ClpA varies linearly with substrate length, $L$, with $R^2 = 1$. Here, $\frac{1}{m} = (0.0352 \pm 0.0001)\ aa^{-1}$, and $n_{intercept} = (-1.44 \pm 0.02)$ steps (errors reported are S.E.).

$k_{obs}$ exhibits a slight increase in $L$, but averaged over triplicates, all three values are within error, see Figure S5 (D). $k_{end}$ is independent of substrate length as shown in Figure S5 (E). Figure S5 (F) shows that triplicate averages of $n$ increase linearly with $L$. The linear increase is described by

$$n = \frac{L}{m} + n_{intercept}, \qquad \text{(S-III)}$$

where $m$ is defined as the kinetic step size and represents the average number of aa processed between two rate-limiting steps.

The analysis was repeated on time courses collected with ClpAP and all three RepA-Titin$_X$ substrates also at 5 mM ATP, see Figure S6 (A)-(C). Individual fits of the time courses reveal that $k_{obs}$ and $k_{end}$, when plotted against $L$ as shown in Figure S6 (D)-(E), respectively, are within error. The $n$ vs. $L$ plot in Figure S5 (F) is linear following Eq. (S-III).

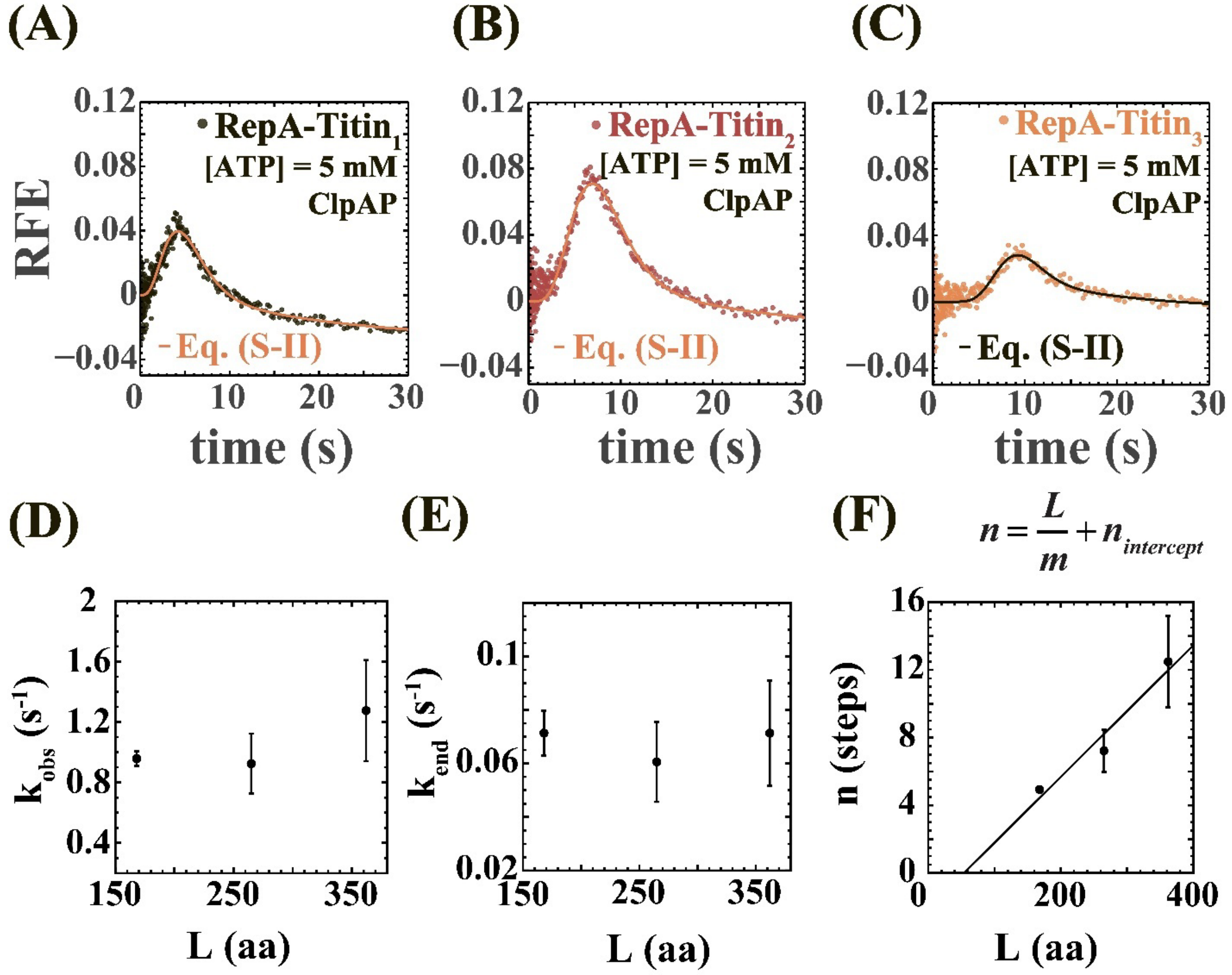


**Figure S6:** **(A)-(C)** Time courses from RepA-Titin$_1$, RepA-Titin$_2$ and RepA-Titin$_3$, respectively, with ClpAP at 5 mM ATP. All time courses were individually fit to Eq. (S-II). **(D)-(E)** $k_{obs}$ and $k_{end}$, respectively, of ClpAP are within error for each substrate. **(L)** $n$ for ClpAP vs. $L$ is linear with $R^2$ = 0.95. Here, $\frac{1}{m}$ = (0.0389 ± 0.0089) aa$^{-1}$, and $n_{intercept}$ = (-2.10 ± 2.45) steps.

In Figure S5 (F) and S6 (F), we see positive $L_{intercept}$, which we interpret as the number of aa that are not part of the unfolding/translocation process. However, we first need to verify if $L_{intercept}$, and thus, $n_{intercept}$ are negligible before drawing any conclusions.

Since $k$ and $k_{end}$ are independent of $L$, and $n$ depends linearly on $L$ following Eq. (S-III), we used Eq. (S-II) and (S-III) to arrive at the following function for fitting time courses from three different substrate lengths simultaneously at each [ATP].

$$RFE(t) = \mathcal{L}^{-1}\left( A_{n-1} \frac{k_{obs}^{\frac{L}{m}+n_{\text{intercept}}-1}}{(k_{obs}+s)^{\frac{L}{m}+n_{\text{intercept}}}} + A_n \frac{k_{obs}^{\frac{L}{m}+n_{\text{intercept}}}}{(k_{end}+s)(k_{obs}+s)^{\frac{L}{m}+n_{\text{intercept}}}} + A_{xU} \frac{k_{end} k_{obs}^{\frac{L}{m}+n_{\text{intercept}}}}{s(k_{end}+s)(k_{obs}+s)^{\frac{L}{m}+n_{\text{intercept}}}} \right)$$

(S-IV)

### S.6. Globally Fitting Time Courses to Confirm the Number of Intermediates

Eq. (S-IV) is derived from Eq. (S-II), which best describes individual time courses. Thus, we expect Eq. (S-IV) to globally describe time courses from ClpA or ClpAP with all three substrates better than Eq. (S-IV) with $A_{n-1} = 0$, i.e. an equation with contributions from one intermediate and the released substrate.

In this analysis, $k_{obs}$, $k_{end}$, and $m$ were constrained to be the same for all three substrates, i.e. global. $A_{n-1}$, $A_n$ and $A_{XU}$ were allowed to be different for each substrate, i.e. local. Figure S7 (A) shows time courses collected using all substrates with ClpA at 5 mM ATP fit globally to Eq. (S-IV) with $A_{n-1} = 0$. Figure S7 (B) shows the same time courses globally fit to Eq. (S-IV). Even though the improvement in fit at 5 mM ATP is visually imperceptible, an $F$-test on the fit $RSS$-values at a level of significance $\alpha = 0.01$ tells us that the fit in Figure S7 (B) is a statistically significant improvement over the fit in Figure S7 (A).

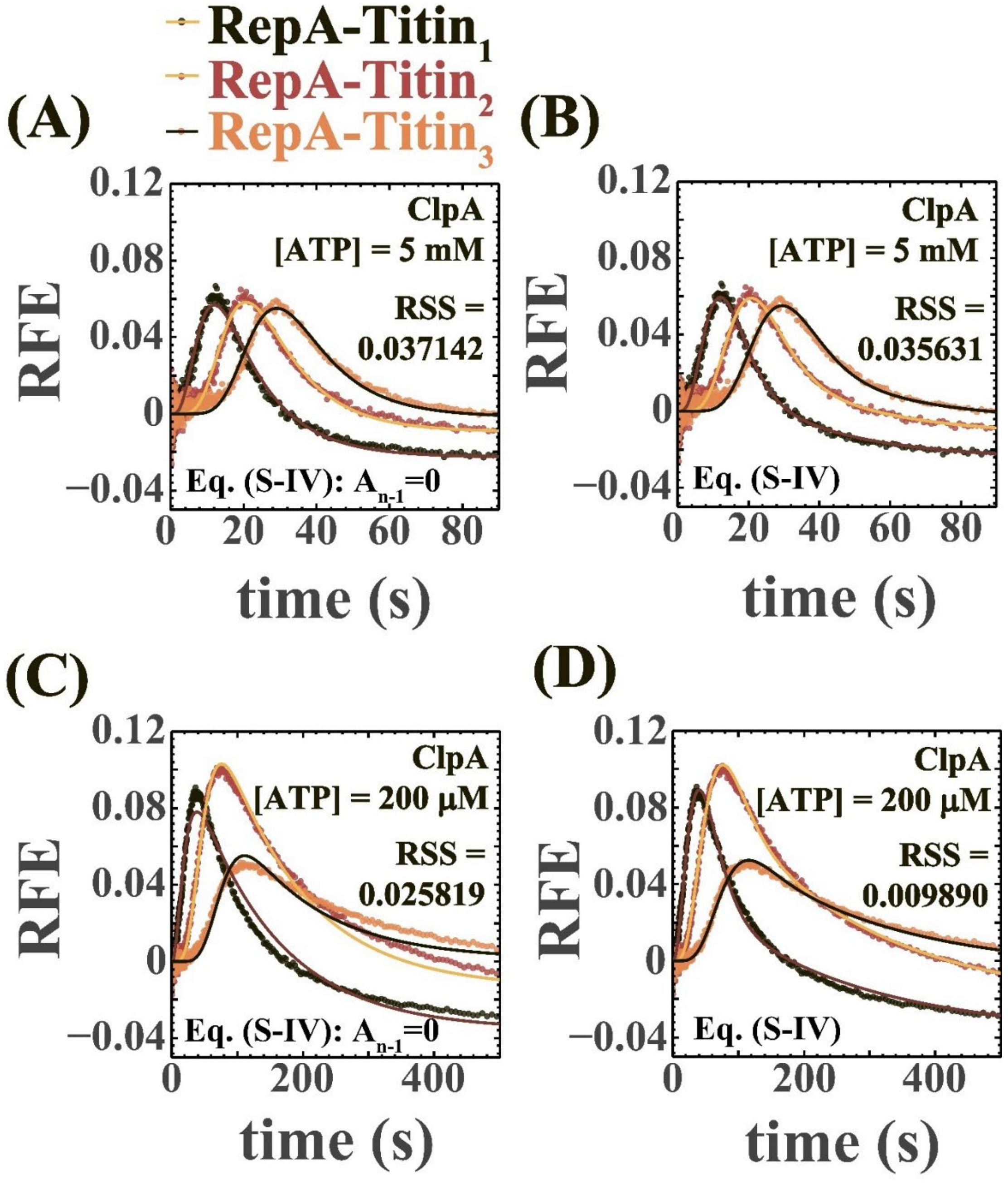


**Figure S7:** **(A)** Time courses from ClpA at high [ATP] globally fit to Eq. (S-IV) with $A_{n-1} = 0$. **(B)** Same time courses from (A) fit to Eq. (S-IV). We do not visually see an improvement with Eq. (S-IV). However, an $F$-test involving the fit $RSS$-values at a level of significance $\alpha = 0.01$ says that (B) is an improvement over (A). **(C)** Time courses from ClpA at low [ATP] globally fit to Eq. (S-IV) with $A_{n-1} = 0$ over all substrates. **(D)** We visually see an improvement when fitting the same time courses from (C) with Eq. (S-IV). An $F$-test involving the fit $RSS$-values at a level of significance $\alpha = 0.01$ confirms that (D) is an improvement over (C).

The analysis was repeated at 200 μM [ATP]. Figure S7 (C) shows data collected using all substrates with ClpA at 200 μM [ATP] fit globally to Eq. (S-IV) with $A_{n-1}$ = 0. Figure S7 (D) shows the same data globally fit to Scheme 1 (Eq. (S-IV)). We visually see an improvement in the fit going from Figure S7 (C) to (D), which is confirmed by an F-test on the fit RSS-values using a level of significance $\alpha = 0.01$.

We carried out similar tests using ClpAP at 5 mM and 200 μM [ATP], see Figure S8. We found that Eq. (S-IV) yields statistically significant improvements in fit compared to Eq. (S-IV) with $A_{n-1} = 0$. In fact, we observe this improvement at all [ATP] for both ClpA and ClpAP. Thus, our decision to include contributions from $I_{n-1}$, $I_n$ and RepA-Titin$_{XU}$ when fitting time courses from three different substrate lengths simultaneously at each [ATP] has been validated.

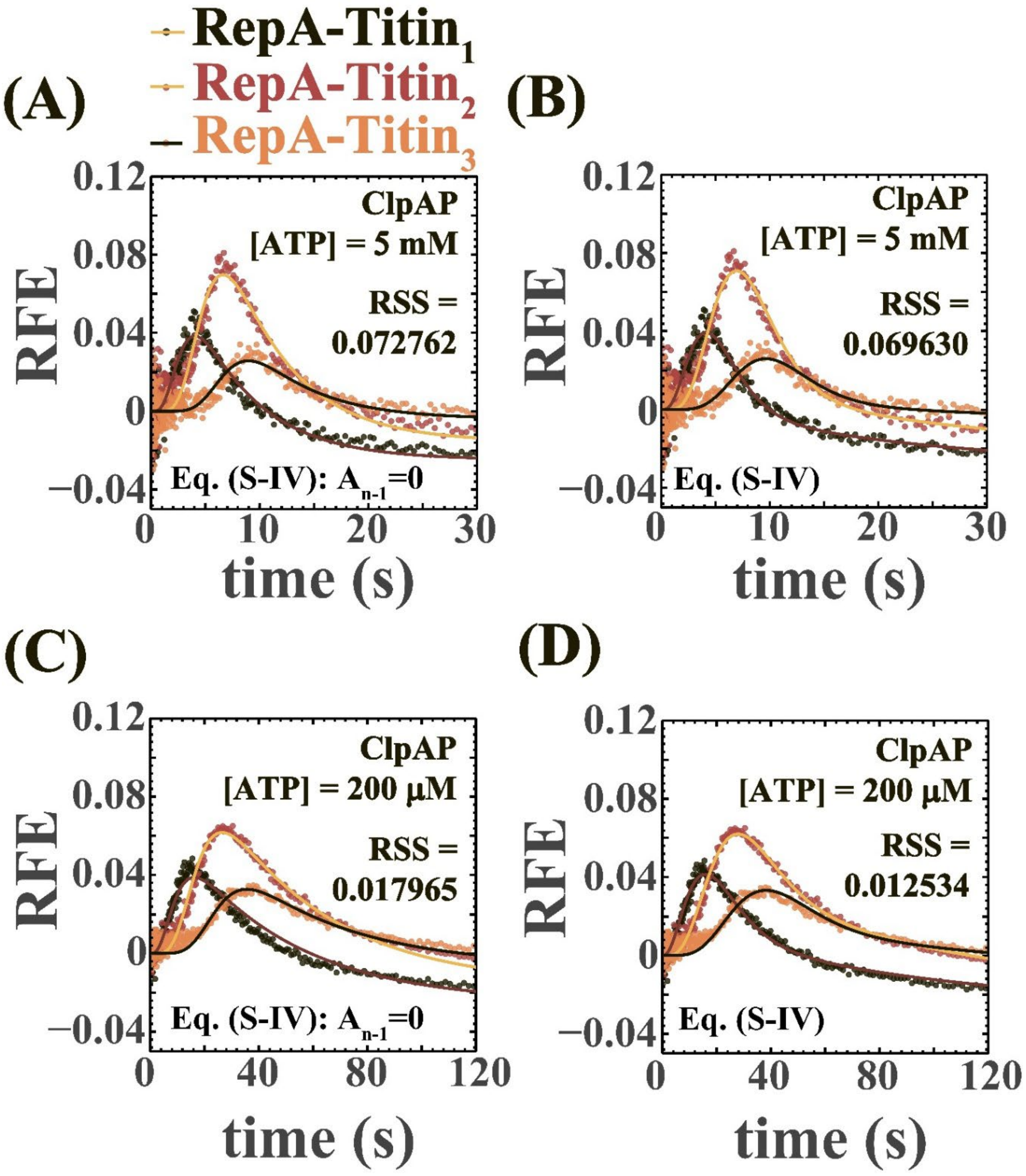


**<u>Figure S8:</u>** **(A)** Time courses from ClpAP at high [ATP] globally fit to Eq. (S-IV) with $A_{n-1} = 0$ over all substrates. **(B)** When we fit the time courses in (A) with Eq. (S-IV), we do not visually see an improvement. However, an *F*-test involving the fit *RSS*-values at a level of significance $\alpha = 0.01$ says that (B) is an improvement over (A). **(C)** Time courses at low [ATP] globally fit to Eq. (S-IV) with $A_{n-1} = 0$ for all substrates. **(D)** We visually see an improvement with Eq. (S-IV), and an *F*-test involving the fit *RSS*-values at a level of significance $\alpha = 0.01$ verifies that (D) is an improvement over (C).

### S.7. Describing Time Courses with the Lowest Number of Optimizable Parameters

Upon fitting time courses from all three substrates simultaneously at each [ATP] using Eq. (S-IV) as shown in Figure S7 (B) and (D) for ClpA, and Figure S8 (B) and (D) for ClpAP, we found that $n_{intercept} < 1$ step. Observing this fraction of a step for $n_{intercept}$, we hypothesized that if the length of the amino acid chain is zero, there should be no steps taken by ClpA or ClpAP. This led us to modify Eq. (S-IV) with $n_{intercept} = 0$ to obtain the following equation.

$$RFE(t) = \mathcal{L}^{-1}\left(A_{n-1}\frac{k_{obs}^{\frac{L}{m}-1}}{(k_{obs}+s)^{\frac{L}{m}}} + A_n\frac{k_{obs}^{\frac{L}{m}}}{(k_{end}+s)(k_{obs}+s)^{\frac{L}{m}}} + A_{xU}\frac{k_{end}k_{obs}^{\frac{L}{m}}}{s(k_{end}+s)(k_{obs}+s)^{\frac{L}{m}}}\right) \quad \text{(S-V)}$$

For both ClpA and ClpAP, Eq. (S-V), embodying Scheme 1, was used to globally fit time courses collected using all substrates at each [ATP]. In this analysis, $k_{obs}$, $k_{end}$, and $m$ were all constrained to be global parameters for all three substrates. $A_{n-1}$, $A_n$ and $A_{XU}$ were local to each substrate. Eq. (S-V) describes the time courses well as shown in Figure S9.

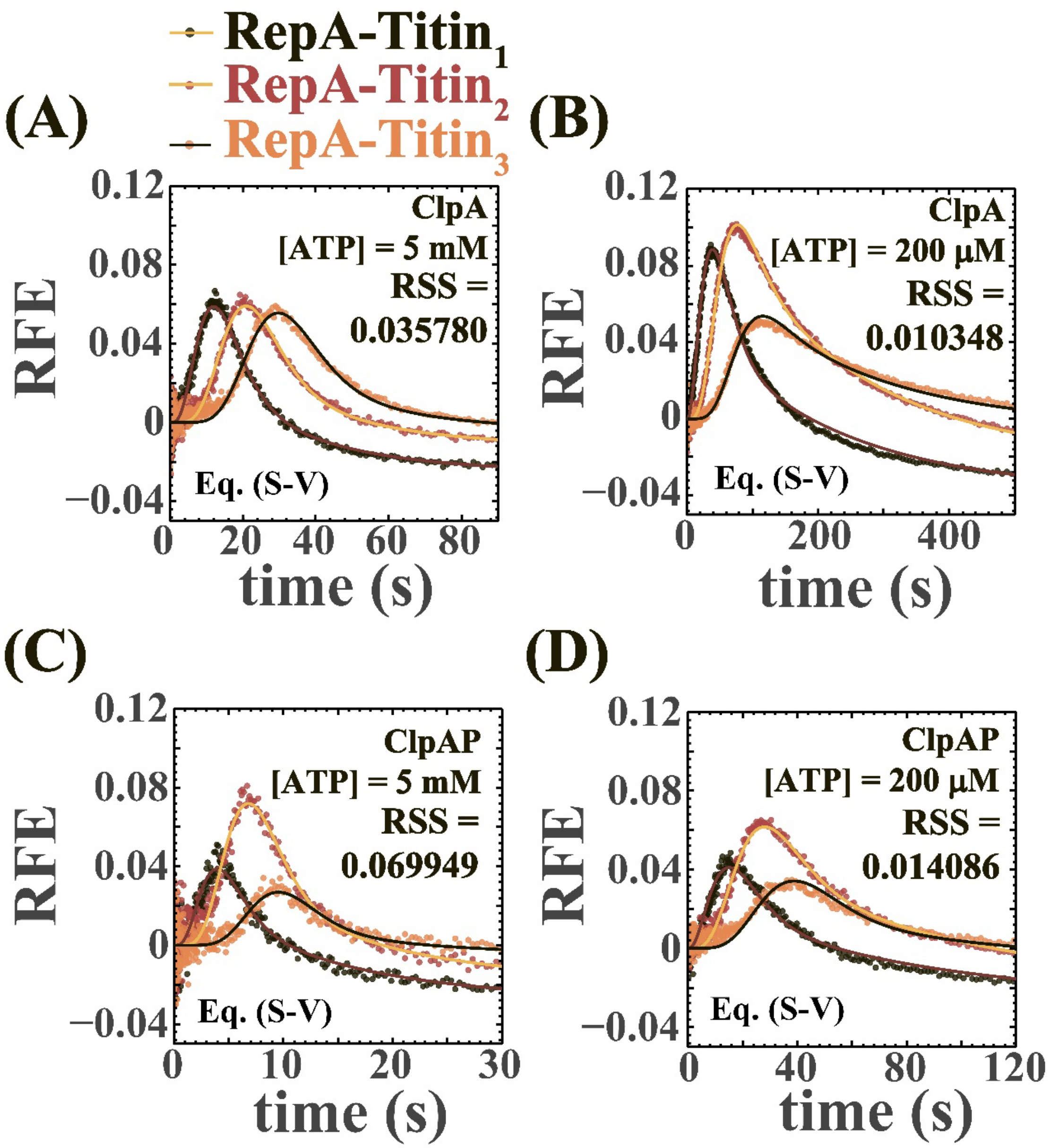


**Figure S9:** **(A)-(D)** Time courses globally fit using Eq. (S-V). *RSS* values have increased compared to Figure S6 (B), (D) and Figure S7 (B), (D). This is expected because we have decreased the number of optimizable parameters in the fits.

Despite the slight increase in fit *RSS*, we accepted the fits because the correction applied to the fitting Eq. (S-V) takes observations from previous fits using Eq. (S-IV) into account.

### S.8. Taking Excluded Length into Account

We interpreted intercepts from $L$ vs. $t_{peak}$ plots shown in Figure S2 as the number of aa from the N-terminus of the substrate bound in the axial channel of ClpA or ClpAP prior to ATP-driven translocation. $L_{intercept}$ was found to be ~18 aa on average, except for ClpA at 200 and 300 μM [ATP]. 18 aa is comparable to the minimum length necessary for ClpA and ClpAP to bind and unfold folded substrates [2].

This is why we assumed that these 18 aa should not be available for ATP-driven translocation. This led us to make a correction to Eq. (S-V) with $L$ being replaced by ($L$-18). Thus, we get the following equation used to fit all time courses.

$$RFE(t) = \mathcal{L}^{-1}\left( A_{n-1} \frac{k_{obs}^{\frac{L-18}{m}-1}}{(k_{obs}+s)^{\frac{L-18}{m}}} + A_n \frac{k_{obs}^{\frac{L-18}{m}}}{(k_{end}+s)(k_{obs}+s)^{\frac{L-18}{m}}} + A_{xU} \frac{k_{end} k_{obs}^{\frac{L-18}{m}}}{s(k_{end}+s)(k_{obs}+s)^{\frac{L-18}{m}}} \right) \tag{S-VI}$$

Representative global fits of time courses to Eq. (S-VI) are shown in Figure S10. The global parameters thus optimized are shown in Table S2.

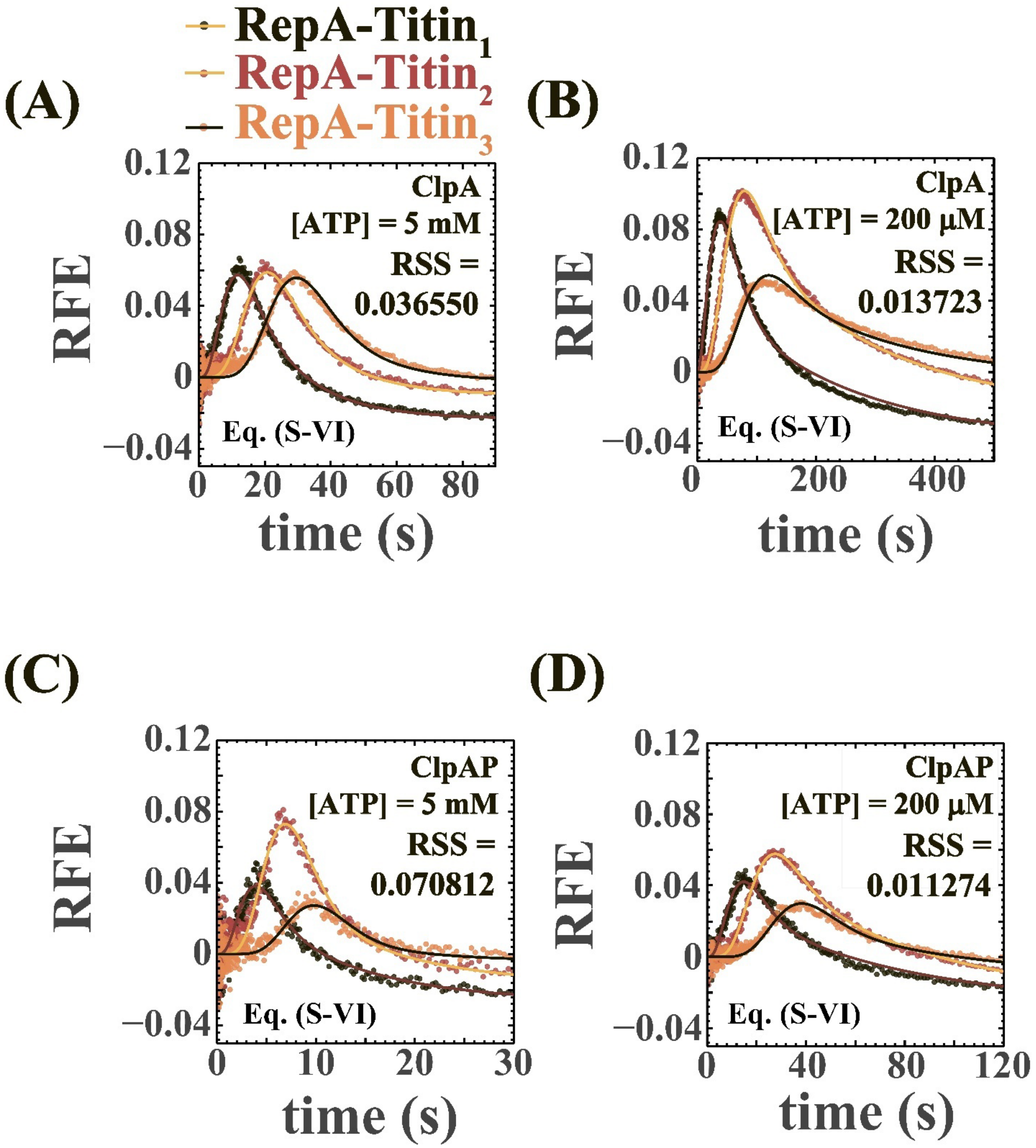


**Figure S10:** **(A)-(D)** Time courses globally fit using Eq. (S-VI). When we compare Figure S9 (A) to Figure S10 (A), Figure S9 (B) to Figure S10 (B), and Figure S9 (C) to Figure S10 (C), we observe the *RSS* to increase slightly. When we compare Figure S9 (D) to Figure S10 (D), we observe the RSS-value to decrease slightly.

Despite the slight increase in fit *RSS* for some fits and slight decrease in others, we accepted the fits in Figure S10 because the correction applied to the fitting Eq. (S-

VI) takes the observation regarding excluded length due to substrate binding from the model-independent peak time analysis into account.

Despite evidence for an excluded length of 18 aa from model-independent peak-time analysis, the time courses are sufficiently described when fit to Eq. (S-V). This is because all substrates are long compared to 18 aa. Our shortest folded substrate is 168 aa long. 18 aa pre-bound in the ClpA axial channel is 11% of 168. This is not a large percentage, allowing us to describe time courses assuming $n = \frac{L}{m}$, with no $L_{intercept}$. Even then, we cannot be oblivious to the evidence for excluded length provided by $L$ vs. $t_{peak}$ plots. So, we decided to correct our fitting function to get Eq. (S-VI).

**Table S2:** Parameters global to all substrate lengths optimized using genetic algorithms and NLLS, shown as a function of [ATP]. Eq. (S-VI) embodying Scheme 1 was used to obtain all parameters except for parameters in columns designated "From Peak Time Analysis". Variables with bars on top are averages over triplicates at each [ATP]. S.D. stands for standard deviations. S.E. stands for standard error.

| **ClpA** | | | | | |
|---|---|---|---|---|---|
| **[ATP] (mM)** | **Repeating Rate Constant, $(\overline{k_{obs}} \pm S.D._k)$ (s$^{-1}$)** | **Dissociation Rate Constant, $(\overline{k_{end}} \pm S.D._{k_{end}})$ (s$^{-1}$)** | **Kinetic Step Size, $(\overline{m_{obs}} \pm S.D._m)$ (aa)** | **Rates (aa s$^{-1}$): $(\overline{m_{obs}k_{obs}} \pm S.D._{mk})$ (aa s$^{-1}$)** | **Rates (aa s$^{-1}$): From Peak Time Analysis, $(v \pm S.E._v)$ (aa s$^{-1}$)** |
| 5 | 0.36 ± 0.02 | 0.052 ± 0.007 | 30.5 ± 1.1 | 11.1 ± 0.2 | 11.6 ± 0.1 |
| 3 | 0.45 ± 0.09 | 0.060 ± 0.017 | 27.9 ± 3.7 | 12.3 ± 0.8 | 13.3 ± 0.1 |
| 2 | 0.45 ± 0.13 | 0.062 ± 0.009 | 27.9 ± 5.5 | 12.1 ± 1.0 | 12.7 ± 0.2 |
| 1 | 0.39 ± 0.08 | 0.053 ± 0.015 | 30.2 ± 7.4 | 11.5 ± 1.4 | 11.3 ± 0.2 |
| 0.75 | 0.35 ± 0.06 | 0.038 ± 0.004 | 28.3 ± 3.3 | 9.7 ± 0.6 | 10.3 ± 0.2 |
| 0.5 | 0.27 ± 0.05 | 0.027 ± 0.006 | 29.7 ± 3.0 | 8.0 ± 0.6 | 8.2 ± 0.0 |
| 0.3 | 0.13 ± 0.05 | 0.010 ± 0.006 | 39.9 ± 9.6 | 4.8 ± 1.2 | 4.6 ± 0.1 |
| 0.2 | 0.05 ± 0.01 | 0.003 ± 0.001 | 52.0 ± 6.6 | 2.5 ± 0.4 | 2.1 ± 0.2 |
| **ClpAP** | | | | | |
| **[ATP] (mM)** | **Repeating Rate Constant, $(\overline{k_{obs}} \pm S.D._k)$ (s$^{-1}$)** | **Dissociation Rate Constant, $(\overline{k_{end}} \pm S.D._{k_{end}})$ (s$^{-1}$)** | **Kinetic Step Size, $(\overline{m_{obs}} \pm S.D._m)$ (aa)** | **Rates (aa s$^{-1}$): $(\overline{m_{obs}k_{obs}} \pm S.D._{mk})$ (aa s$^{-1}$)** | **Rates (aa s$^{-1}$): From Peak Time Analysis, $(v \pm S.E._v)$ (aa s$^{-1}$)** |
| 5 | 1.10 ± 0.14 | 0.085 ± 0.008 | 29.7 ± 2.9 | 32.3 ± 1.0 | 37.7 ± 0.4 |
| 3 | 1.35 ± 0.21 | 0.102 ± 0.002 | 24.4 ± 3.3 | 32.4 ± 0.8 | 33.7 ± 0.4 |
| 2 | 1.06 ± 0.18 | 0.078 ± 0.013 | 27.3 ± 2.5 | 28.5 ± 2.0 | 31.9 ± 0.5 |
| 1 | 0.91 ± 0.08 | 0.075 ± 0.012 | 27.6 ± 1.9 | 25.1 ± 1.5 | 27.7 ± 0.5 |
| 0.75 | 0.82 ± 0.10 | 0.076 ± 0.003 | 29.6 ± 3.6 | 24.1 ± 0.8 | 24.5 ± 1.3 |
| 0.5 | 0.56 ± 0.07 | 0.049 ± 0.002 | 32.2 ± 2.6 | 17.8 ± 1.0 | 20.8 ± 1.0 |
| 0.3 | 0.34 ± 0.05 | 0.027 ± 0.001 | 35.9 ± 4.9 | 12.0 ± 0.4 | 15.1 ± 0.3 |
| 0.2 | 0.20 ± 0.05 | 0.013 ± 0.002 | 40.4 ± 5.5 | 8.0 ± 0.7 | 9.0 ± 0.5 |

### S.9. Probing Cooperativity with Respect to [ATP] for ClpA and ClpAP using Parameters from Global Fits

Figure S11 shows the parameters displayed in Table S2 and Figure 5, fit to the cooperative (solid lines) and noncooperative (dashed lines) models. $F$-tests were carried out following the method detailed in Sup. Section S.2 with $\alpha = 0.01$ for Figure S10 (A), (C), (D), and (F). For Figure S10 (B), and (E) we used $\alpha = 0.05$. Decisions regarding the models best describing the trends are given in the figure legend and in Table S3. Also, the fit parameters best describing the trends are given in Table S3.

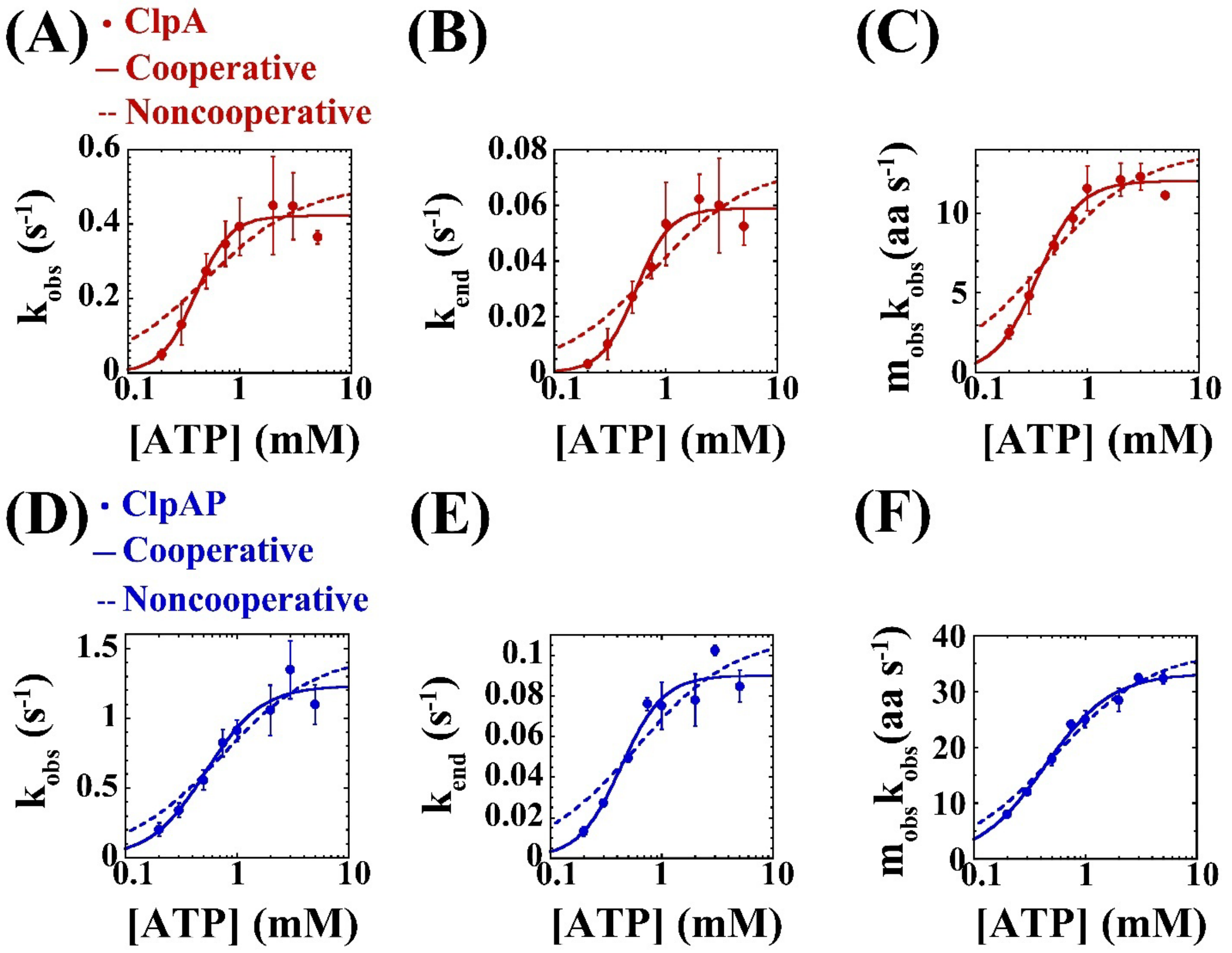


**Figure S11:** Plots of **(A)** $k_{obs}$ as a function of [ATP] for ClpA in Figure 5 (C); **(B)** $k_{end}$ as a function of [ATP] for ClpA in Figure 5 (D); **(C)** $m_{obs}k_{obs}$ as a function of [ATP] for ClpA in Figure 5 (E); **(D)** $k_{obs}$ as a function of [ATP] for ClpAP in Figure 5 (C); **(E)** $k_{end}$ as a function of [ATP] for ClpAP in Figure 5 (D); and **(F)** $m_{obs}k_{obs}$ as a function of [ATP] for ClpAP in Figure 5 (E). All plots were fit to both the cooperative (solid lines) and noncooperative models (dashed lines). **(A)-(F)** Error bars represent S.D. **(A), (C), (D) and (F)** $F$-tests with $\alpha = 0.01$ support the cooperative model for ClpA and the noncooperative model for ClpAP. **(B) and (E)** Both plots visually appear to saturate within one log unit. Because of this, we trusted less stringent $F$-tests with $\alpha = 0.05$ to decide that both trends follow the cooperative model.

**Table S3:** Models and fit parameters best describing trends in Figure S11 and Figure 5 of the main paper. All errors represent S.E.

| **ClpA** | | | | | | | |
|---|---|---|---|---|---|---|---|
| **Parameter vs. [ATP]** | **Model** | **No. of fit parameters, *p*** | $R^2$ | ***RSS*** | **Parameter$_{max}$** | $K_d$ **(mM)** | ***h*** |
| $k_{obs}$ ($s^{-1}$) | Cooperative | 3 | 0.965 | 0.005 | 0.42 ± 0.02 | 0.40 ± 0.04 | 2.8 ± 0.6 |
| $k_{end}$ ($s^{-1}$) | Cooperative | 3 | 0.974 | 0.00010 | 0.059 ± 0.003 | 0.54 ± 0.05 | 2.9 ± 0.6 |
| $m_{obs}k_{obs}$ (aa $s^{-1}$) | Cooperative | 3 | 0.983 | 1.53 | 12.0 ± 0.4 | 0.36 ± 0.02 | 2.3 ± 0.3 |
| **ClpAP** | | | | | | | |
| **Parameter vs. [ATP]** | **Model** | **No. of fit parameters, *p*** | $R^2$ | ***RSS*** | **Parameter$_{max}$** | $K_d$ **(mM)** | ***h*** |
| $k_{obs}$ ($s^{-1}$) | Noncooperative | 2 | 0.915 | 0.093 | 1.46 ± 0.15 | 0.74 ± 0.22 | 1 (constrained) |
| $k_{end}$ ($s^{-1}$) | Cooperative | 3 | 0.987 | 0.00034 | 0.090 ± 0.006 | 0.43 ± 0.05 | 2.3 ± 0.6 |
| $m_{obs}k_{obs}$ (aa $s^{-1}$) | Noncooperative | 2 | 0.991 | 6.04 | 40.0 ± 1.1 | 0.59 ± 0.05 | 1 (constrained) |

We thus obtained parameters that can be directly compared with parameters obtained in our previous studies using unstructured substrates [3, 4], see Table S4.

**Table S4:** Comparable parameters for ClpA and ClpAP on folded and unstructured substrates. All errors reported are S.E.

| **Motor** | **Folded Substrates** | | | **Unstructured Substrates** | |
|---|---|---|---|---|---|
| | $v_{max}$ (aa $s^{-1}$) | $(m_{obs}k_{obs})_{max}$ (aa $s^{-1}$) | | $(m_Tk_T)_{max}$ (aa $s^{-1}$) | |
| **ClpA** | 12.6 ± 0.4 | 12.0 ± 0.4 | | 19.5 ± 0.7 [3] | |
| **ClpAP** | 41.0 ± 1.2 | 40.0 ± 1.1 | | 36.1 ± 0.7 [4] | |
| | $(k_{obs})_{max}$ ($s^{-1}$) | | | $(k_T)_{max}$ ($s^{-1}$) | |
| **ClpA** | 0.42 ± 0.02 | | | 1.44 ± 0.06 [3] | |
| **ClpAP** | 1.46 ± 0.15 | | | 7.9 ± 0.2 [4] | |
| | $K_d$ **(mM)** | | | $K_d$ **(mM)** | |
| | **From** $k_{obs}$ **vs. [ATP]** | **From** $m_{obs}k_{obs}$ **vs. [ATP]** | **From** $v$ **vs. [ATP]** | **From** $k_T$ **vs. [ATP]** | **From** $m_Tk_T$ **vs. [ATP]** |
| **ClpA** | 0.40 ± 0.04 | 0.36 ± 0.02 | 0.39 ± 0.02 | ~0.56 [3] | ~0.53 [3] |
| **ClpAP** | 0.74 ± 0.22 | 0.59 ± 0.05 | 0.53 ± 0.05 | ~0.21 [4] | ~0.21 [4] |
| | ***h*** | | | ***h*** | |
| | **From** $k_{obs}$ **vs. [ATP]** | **From** $m_{obs}k_{obs}$ **vs. [ATP]** | **From** $v$ **vs. [ATP]** | **From** $k_T$ **vs. [ATP]** | **From** $m_Tk_T$ **vs. [ATP]** |
| **ClpA** | 2.8 ± 0.6 | 2.3 ± 0.3 | 2.4 ± 0.3 | 2.2 ± 0.3 [3] | 2.5 ± 0.3 [3] |
| **ClpAP** | 1 (constrained) | 1 (constrained) | 1 (constrained) | 1 (constrained) [4] | 1 (constrained) [4] |

### S.10. Differential Coupling for ClpAP

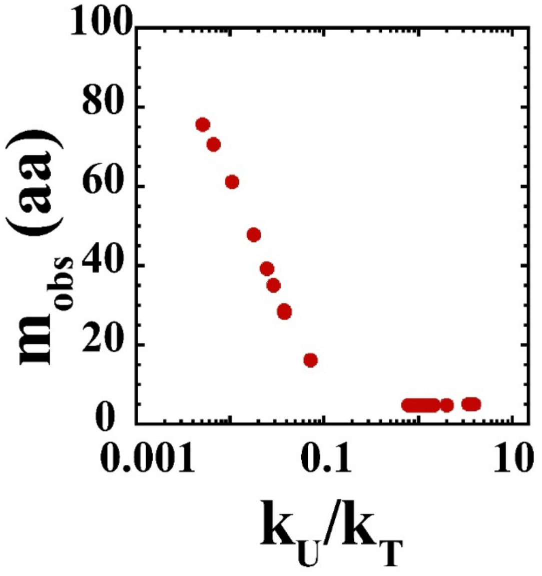


**Figure S12:** Scheme 2 was used to simulate time courses using $m_T$ = 5 aa, $m_U$ = 97 aa, and different values for the translocation rate constant, $k_T$, and the unfolding rate constant, $k_U$. Upon fitting the simulated time courses using Scheme 1, the predicted kinetic step size, $m_{obs}$, was found to vary with $k_U/k_T$ as shown.

### S.10. Comparing Parameters Obtained Using Unstructured and Folded Substrates

Our experiments and analyses yielded parameters that can be directly compared with parameters obtained in our previous studies using unstructured substrates [3, 4].

**Table S4:** Comparable parameters for ClpA and ClpAP on folded and unstructured substrates. All errors reported are S.E.

| Motor | Folded Substrates | | | Unstructured Substrates | |
|---|---|---|---|---|---|
| | $v_{max}$ (aa $s^{-1}$) | $(m_{obs}k_{obs})_{max}$ (aa $s^{-1}$) | | $(m_Tk_T)_{max}$ (aa $s^{-1}$) | |
| **ClpA** | 12.6 ± 0.4 | 12.0 ± 0.4 | | 19.5 ± 0.7 [3] | |
| **ClpAP** | 41.0 ± 1.2 | 40.0 ± 1.1 | | 36.1 ± 0.7 [4] | |
| | $(k_{obs})_{max}$ ($s^{-1}$) | | | $(k_T)_{max}$ ($s^{-1}$) | |
| **ClpA** | 0.42 ± 0.02 | | | 1.44 ± 0.06 [3] | |
| **ClpAP** | 1.46 ± 0.15 | | | 7.9 ± 0.2 [4] | |
| | $K_d$ (mM) | | | $K_d$ (mM) | |
| | **From $k_{obs}$ vs. [ATP]** | **From $m_{obs}k_{obs}$ vs. [ATP]** | **From $v$ vs. [ATP]** | **From $k_T$ vs. [ATP]** | **From $m_Tk_T$ vs. [ATP]** |
| **ClpA** | 0.40 ± 0.04 | 0.36 ± 0.02 | 0.39 ± 0.02 | ~0.56 [3] | ~0.53 [3] |
| **ClpAP** | 0.74 ± 0.22 | 0.59 ± 0.05 | 0.53 ± 0.05 | ~0.21 [4] | ~0.21 [4] |
| | $h$ | | | $h$ | |
| | **From $k_{obs}$ vs. [ATP]** | **From $m_{obs}k_{obs}$ vs. [ATP]** | **From $v$ vs. [ATP]** | **From $k_T$ vs. [ATP]** | **From $m_Tk_T$ vs. [ATP]** |
| **ClpA** | 2.8 ± 0.6 | 2.3 ± 0.3 | 2.4 ± 0.3 | 2.2 ± 0.3 [3] | 2.5 ± 0.3 [3] |
| **ClpAP** | 1 (constrained) | 1 (constrained) | 1 (constrained) | 1 (constrained) [4] | 1 (constrained) [4] |

### S.12. References


1. Makridakis, S., S.C. Wheelwright, and R.J. Hyndman, *FORECASTING METHODS AND APPLICATIONS, 3RD ED*. 2008: Wiley India Pvt. Limited.
2. Hoskins, J.R., S.Y. Kim, and S. Wickner, *Substrate recognition by the ClpA chaperone component of ClpAP protease.* J Biol Chem, 2000. **275**(45): p. 35361-7.
3. Rajendar, B. and A.L. Lucius, *Molecular mechanism of polypeptide translocation catalyzed by the Escherichia coli ClpA protein translocase.* J Mol Biol, 2010. **399**(5): p. 665-79.
4. Miller, J.M., et al., *E. coli ClpA catalyzed polypeptide translocation is allosterically controlled by the protease ClpP.* J Mol Biol, 2013. **425**(15): p. 2795-812.
